# A protein hydroxylase couples epithelial membrane biology to nucleolar ribosome biogenesis

**DOI:** 10.1101/2023.03.15.532818

**Authors:** Eline Hendrix, Regina Andrijes, Uncaar Boora, Arashpreet Kaur, James R Bundred, Leah Officer-Jones, Rachel Pennie, Ian R Powley, Adam Zayer, Raphael Heilig, Christian A E Westrip, Sally C Fletcher, Charlotte D Eaton, Tristan J Kennedy, Sonia Piasecka, Roman Fischer, Stephen J Smerdon, John Le Quesne, Mathew L Coleman

## Abstract

Jumonji-C (JmjC) ribosomal protein hydroxylases are an ancient class of oxygen- and Fe(II)-dependent oxygenases that spawned the wider JmjC family and Histone Lysine Demethylases (KDMs) in eukaryotes. Myc-induced Antigen (MINA) has been implicated in ribosome biogenesis and was assigned as a nucleolar-localized JmjC histidyl hydroxylase of the large ribosomal subunit protein RPL27A, consistent with reports that it supports cell growth and viability in a variety of tumor cell types. Reported roles in diverse aspects of disease biology may be consistent with additional MINA functions, although the molecular mechanisms involved remain unclear. Here, we describe an extra-nucleolar interaction of MINA with the Hinge domain of the membrane-associated guanylate kinase, MPP6. We show that MINA promotes the expression and membrane localization of MPP6 and that the MINA-MPP6 pathway is required for epithelial tight junction integrity and barrier function. The function of MINA in this novel pathway is suppressed by ribosomal RNA transcription and the nucleolar MINA interactome. In this way, MINA couples epithelial membrane biology to nucleolar ribosome biogenesis. Our work sheds light on how quiescent cells lose adhesion as they switch to proliferative states associated with increased ribosome biogenesis.

## Introduction

Cross-talk between cellular processes is key to enabling a coordinated multisystem response to a change in cell fate. The switch to a proliferative state requires increased metabolic activity and accumulation of biomass to feed cell growth and cell cycle-dependent changes in adhesion to support division^1–3^. Cells have evolved sophisticated signaling mechanisms to couple such processes. For example, bidirectional cross-talk has been demonstrated between cell adhesion and the cell cycle^3^. Whether cell adhesion is coupled to other fundamental processes linked to growth remains unclear.

Ribosomal RNA (rRNA) synthesis is a rate-limiting step of ribosome biogenesis and is upregulated as cells switch from growth-arrested to proliferative states in response to major growth responsive signaling pathways and transcription factors, such as Myc^2, 4^. Activation of Myc or the RNA polymerase I (RNApol1) transcription initiation factor RRN3 is sufficient to induce RNApol1 activity and proliferation in human epithelial cells^5–8^. Remarkably, normal 3D morphogenesis (which requires functional polarity and adhesion) is impaired under such conditions^7, 8^, suggesting that ribosome biogenesis, like the cell cycle, might also be coupled to epithelial membrane biology in some way.

MINA is a nucleolar-localized Myc target gene implicated in ribosome biogenesis^9–11^. It is a member of the 2-oxoglutarate (2OG)-, Fe(II)-, and oxygen-dependent oxygenase family, and is highly related to NO66, a prokaryotic orthologue of which is believed to have evolved into the eukaryotic JmjC enzymes, including KDMs^12^. Despite the presence of a JmjC catalytic domain, detailed structural and biochemical studies suggest that MINA and NO66 may not be KDMs^12^. Indeed, they catalyze non-redundant and site-specific histidine hydroxylation of the large ribosomal subunit proteins RPL27A and RPL8, respectively^13^. Hydroxylation has been proposed to support the optimal conformation of the target residue to promote its interaction with rRNA^14^. Consistent with a positive role in ribosome biogenesis/function, multiple studies have described upregulation of MINA in rapidly growing cells, including in tumors (reviewed in^11^). However, recent reports have also implicated MINA in suppressing cell migration, invasion, and epithelial-to-mesenchymal transition (EMT)^15, 16^. How MINA controls these aspects of epithelial biology, and whether it is independent of, or may be coupled to, its role in nucleolar ribosome biogenesis, remains unclear.

Here, we describe a novel extra-nucleolar function for MINA in epithelial barrier function through its pseudosubstrate-like interaction with MPP6, a member of the ‘palmitoylated membrane protein’ (MPP) subfamily of the MAGUK superfamily of cell adhesion and polarity adaptor proteins. Furthermore, we provide evidence suggesting that increased RNAPol1 activity sinks MINA into the nucleolus to suppress the role of MINA in cell adhesion, thereby coupling ribosome biogenesis to epithelial membrane biology.

## Results

### MINA promotes epithelial barrier function

Although MINA has been implicated as an oncogenic driver of tumor proliferation, there is emerging literature on its role in suppressing cell migration and invasion. However, these functional studies have generally focused on the role of MINA in highly proliferative cells that have undergone a robust EMT, such as rapidly proliferating tumor cell lines^11^ and references therein)). MINA is ubiquitously expressed in normal healthy human tissues (Figure S1), and yet its role in such contexts is poorly characterized. To begin to explore its physiological role in more detail we studied MINA in intestinal epithelial cells that retain good cell adhesion, polarity, and barrier function, and that can differentiate and undergo morphogenesis in 3D^17, 18^. As such, we generated Caco-2 cell lines stably expressing doxycycline-inducible MINA shRNA sequences. In these cells MINA depletion did not affect proliferation (Figure S2) or grossly alter apico-basal asymmetry, as assessed by the localization of tight and adherens junction markers ZO-1 and E-cadherin, respectively (Figure 1A). However, we observed an increase in cell perimeter and cell flattening following MINA knockdown (Figures 1A and 1B). MINA depletion also resulted in delayed formation of transepithelial electrical resistance (TEER; Figure 1C) and a reduction in ‘domes’ (Figures 1D and 1E), which are normally formed when fluid is pumped under a cell monolayer, in the context of good cell:cell adhesion. Taken together, these data suggest that MINA may be required for tight junction (TJ) integrity and barrier function^19, 20^. Consistent with this hypothesis, we noted reduced membrane localization and expression of tight junction-associated protein Claudin-1 in response to MINA depletion (Figures 1F, 1G and 1H). How a nucleolar ribosomal protein hydroxylase regulates such membrane-related biology is unclear.

**Figure 1.**
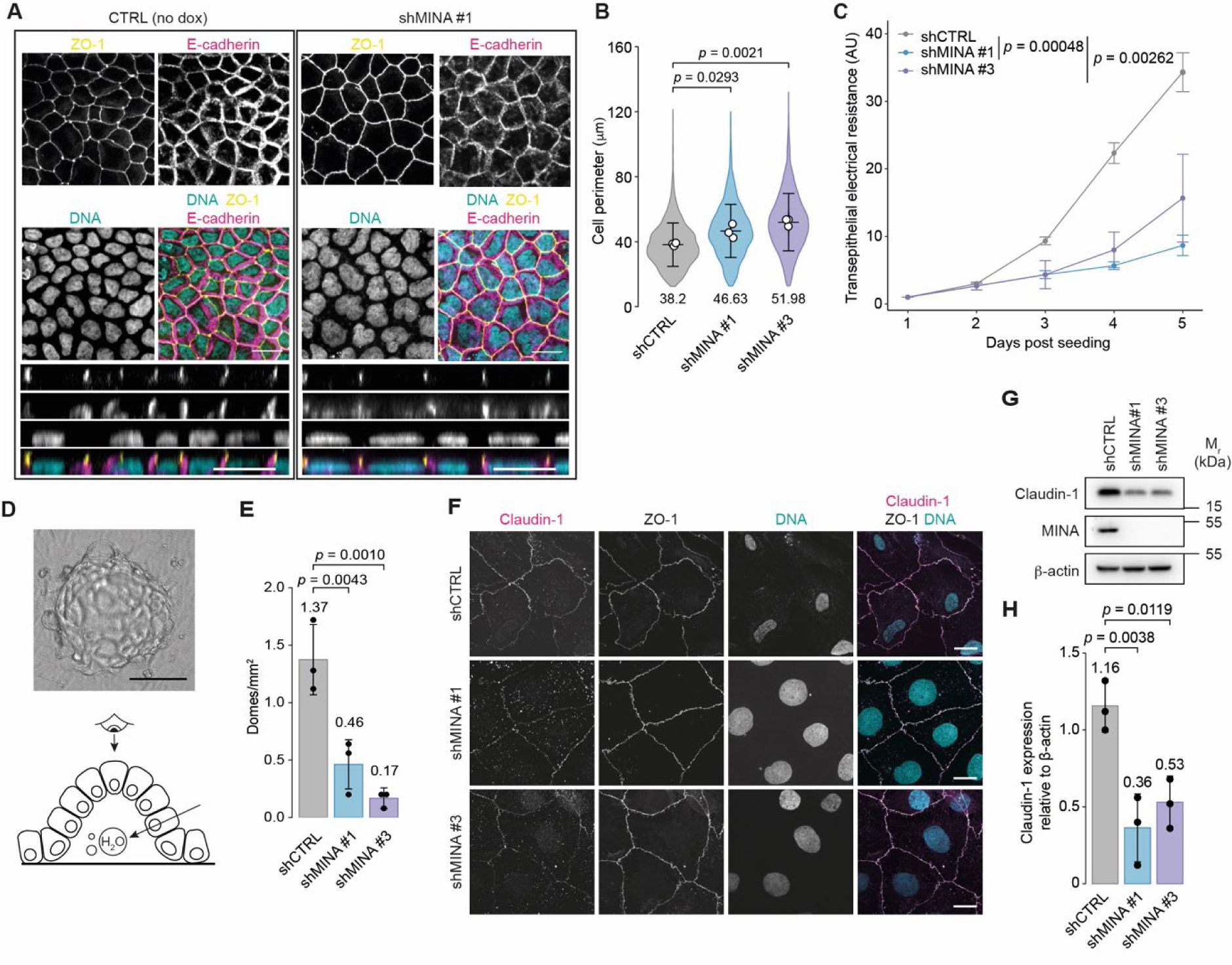
MINA promotes tight junction integrity and epithelial barrier function. **(A)** Increased perimeter of cells in MINA-depleted monolayers. Caco-2 cells expressing a non-targeting (shCTRL) or MINA-targeting shRNA (shMINA#1) were grown on a semi-permeable membrane for 10 days and stained for ZO-1 (yellow), E-cadherin (magenta) and DNA (DAPI, cyan). Consecutive images were taken from the basal to the apical side and presented as a maximum projection image (top panels) or orthogonal view (rectangular bottom panels). Orthogonal views show the basal side at the bottom and apical side (as indicated by ZO-1) at the top. Scale bars, 20 µm. **(B)** Quantification of cell perimeters under conditions presented in (A). Violin plot represents the data from 3 independent experiments. n = 4572 (shCTRL), 2616 (shMINA #1) and 2426 (shMINA #3) cells. The average of each condition is shown underneath each respective violin. **(C)** Delayed establishment of transepithelial electrical resistance (TEER) in MINA-depleted cells. Monolayer integrity and permeability were analyzed on the indicated days by measuring the electrical resistance across the monolayer. Line graph shows the average of three technical repeats of one representative biological experiment. The experiment was repeated 5 times with similar results. **(D)** Top, phase-contrast image of a dome, as viewed from the top. Scale bar, 100 µm. Bottom, illustration of a dome, as viewed from the side. **(E)** Quantification of dome formation in monolayers of stable Caco-2 cells expressing the indicated shRNA sequences. Bars represent the average number of domes per mm^2^ of n = 3 independent experiments. The average of each condition is shown above each respective bar. **(F)** Altered localization of tight junction protein Claudin-1 following MINA knockdown. Representative images of Caco-2 monolayers expressing the indicated shRNAs were stained for Claudin (magenta), ZO-1 (gray) and DNA (DAPI; cyan). Scale bars, 20 µm. **(G)** Reduced Claudin-1 protein expression in response to MINA knockdown. Extracts from Caco-2 cells expressing the indicated shRNA sequences were grown into a monolayer over 5 days and immunoblotted for Claudin-1 (23 kDa) and MINA (53 kDa). β-actin was used as loading control. Uncropped images are shown in source data. **(H)** Quantification of (G). Bar graph shows Claudin-1 expression relative to β-actin. Bars represent the average of n = 3 biological repeats. (B), (C), (E), and (H); Statistical significance was determined using a one-way analysis of variance (ANOVA) and p values were calculated post-hoc using a Dunnett’s test for multiple comparisons. Error bars, standard deviation (SD).

### MINA nucleolar localization is dependent on its nucleolar interactome and elevated ribosome biogenesis

To investigate the role of MINA in epithelial membrane biology we first re-evaluated its subcellular localization. Consistent with reports describing nucleolar MINA staining in highly proliferative normal tissues, such as the testes^21^, and in tumors (reviewed in^11^), we observed strong nucleolar enrichment of MINA in rapidly growing mesenchymal-like cell lines (e.g. SW620 and A549; Figure 2A). Surprisingly, this enrichment was diminished in slower growing cell lines with a more epithelial morphology. In U2OS and Caco-2 cells the localization of MINA appeared heterogenous with some cells showing diffuse nucleoplasmic staining and no nucleolar enrichment (Figures 2A and S3A).

**Figure 2.**
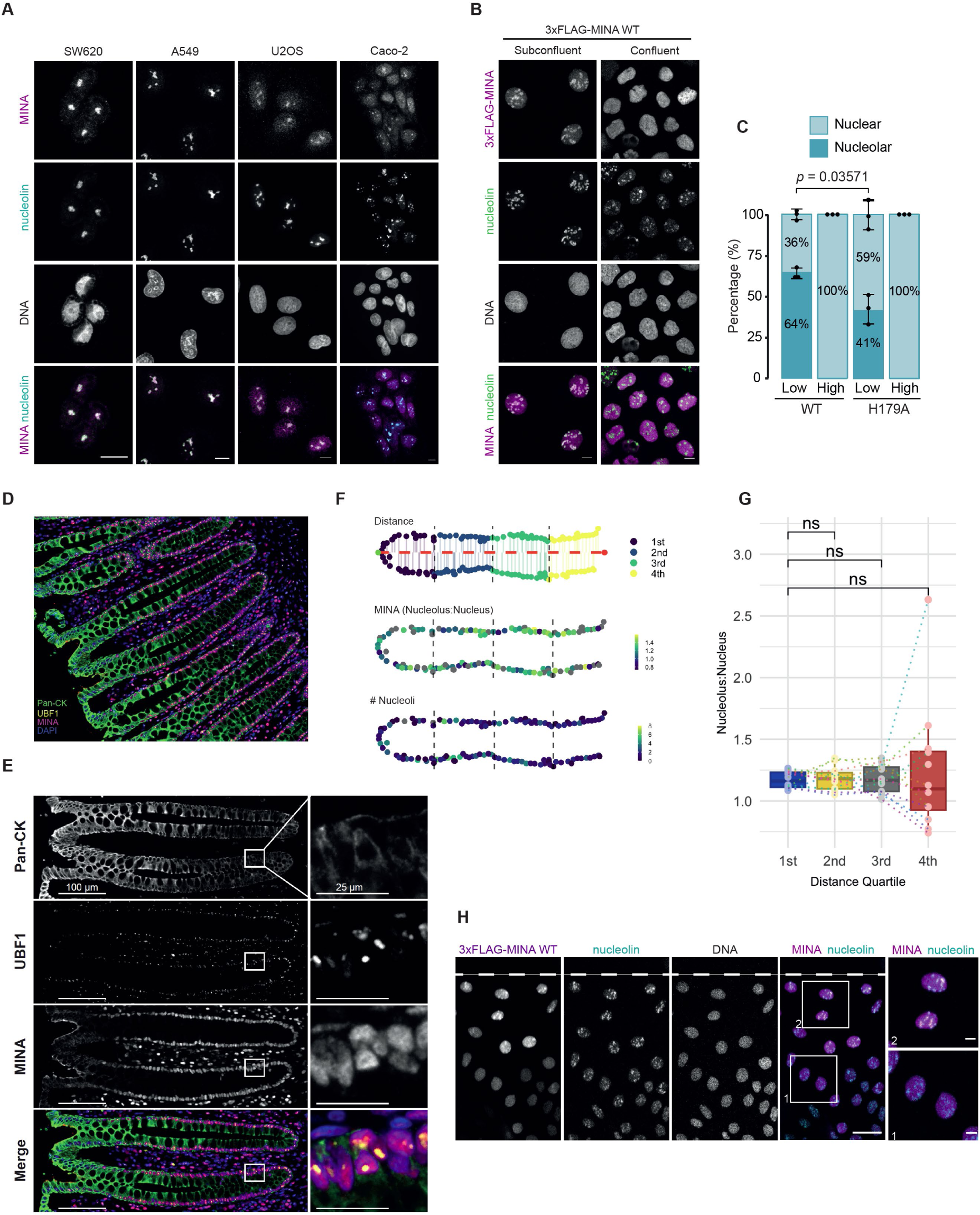
MINA nucleolar localization is regulated by growth and hydroxylase activity. **(A)** Localization of MINA in selected cancer cell lines. Representative immunofluorescence images showing nucleolar and/or nucleoplasmic MINA. Cells were fixed with methanol and stained for endogenous MINA (magenta) and the nucleolar marker nucleolin (cyan). Scale bars, 10 µm. **(B)** Nucleolar MINA localization depends on cell confluence. Representative immunofluorescence images showing the localization of wildtype (WT) 3xFLAG-MINA in subconfluent and confluent conditions. Cells were incubated with 0.1 µg/mL doxycycline for 48 h to induce expression of 3xFLAG-MINA and stained with an antibody raised against endogenous MINA (magenta). To confirm that the detected signal was of overexpressed, exogenous 3xFLAG-MINA, an empty vector control was used (Figure S3B). Cells were counterstained for nucleolin (NCL, green) and DNA (DAPI, gray). Scale bars, 10 µm **(C)** Bar graph showing the percentage of cell in which nucleolar enrichment could (‘Nucleolar’) or could not (‘Nuclear’) be observed. ‘Low’ and ‘High’ refer to the level of confluence. Bars show the average of three biological repeats. Error bars, SD. A total of one hundred cells were counted per condition per experiment. Samples were blinded before analysis. Statistical significance was determined using an unpaired t-test. **(D)** Localization of MINA in normal human colon. Representative multiplexed immunofluorescence image showing MINA (magenta), epithelial cell marker pan-cytokeratin (green) and the nucleolar marker UBF1 (yellow). **(E)** Representative immunofluorescence images showing the localization of MINA, pan-cytokeratin and the nucleolar marker UBF1 in normal human colonic crypts. Enlargements show nuclear detail with pan-nuclear MINA localization. Note the loss of MINA staining intensity in the differentiated cells of the villi. **(F)** Nucleolar number, nucleoplasm and nucleolar MINA expression were quantified at the cellular level (n = 12 crypts) and crypts divided into quartiles based on their distance from the crypt base. Nucleolus:Nuclear ratio calculated as mean nucleolar per pixel intensity divided by mean nucleoplasm per pixel intensity. Nucleoli # represents number of nucleolar quantified per cell nucleus. **(G)** Per cell Nucleolus:nuclear MINA expression were calculated as in (F) and binned per quartile along the crypt axis. Mean values were calculated per crypt (n=12, represented as points). Box plots represent median values with hinges corresponding to the first and third quartiles and whiskers extending to no further than 1.5* the interquartile range from the hinge. ns represents non-significant (p>0.05) difference from paired Wilcoxon test. **(H)** MINA becomes enriched in the nucleolus following disruption of a confluent monolayer. A scratch was introduced to disrupt a confluent monolayer of Caco-2 cells expressing wildtype 3xFLAG-MINA. Cells were fixed 24 h after the scratch was made. Re-localization of 3xFLAG-MINA was observed in cells proximal to the scratch (top, indicated by the white dashed line). In contrast, at a distance from the scratch, 3xFLAG-MINA remained in the nucleoplasm (bottom). Scale bars, 50 or 10 (enlargements) µm

We hypothesized that the subcellular localization of MINA might be related to growth status. To test this, we cultured Caco-2 to either subconfluence or confluence (Figure 2B and 2C): In subconfluent cells we observed a more homogenous subcellular distribution of MINA with diffuse nucleoplasmic staining *and* nucleolar enrichment. In confluent cells MINA distributed evenly throughout the nucleus without any nucleolar enrichment (Figure 2B and 2C). To explore the physiological relevance of pan-nuclear MINA localization we developed an automated and quantitative multiplex immunofluorescence assay for staining MINA and nucleoli in normal human colon (Figure 2D). Consistent with our observations in confluent Caco-2 cells, we observed pan-nuclear MINA staining across intestinal crypts and villi (Figure 2E). Quantitative analyses using nuclear (DAPI) and nucleolar (UBF1) markers indicated that the nucleolar:nuclear ratio of MINA staining intensity was close to 1 (Figure 2F and 2G), confirming pan-nuclear localization in healthy intestinal epithelium.

To explore whether MINA localization is dynamic and responds to external cues we grew Caco-2 cells to confluence before scratching the monolayer. As expected, confluent cells distal to the scratch showed pan-nuclear MINA localization. However, cells proximal to the scratch showed elevated levels of MINA and nucleolar enrichment (Figure 2G). The nucleolar enrichment of overexpressed MINA was partly dependent on its catalytic, suggesting a role for RPL27A hydroxylation (Figures 2C and S3B).

Considering the importance of MINA hydroxylase activity, and the established links between growth and ribosome biogenesis, we hypothesized that high levels of ribosome biogenesis might act as a nucleolar sink for MINA. Consistent with this possibility, we observed significant nucleolar enrichment of nascent proteins that colocalized with MINA in subconfluent cells (Figure 3A). Under these conditions, inhibition of rRNA transcription and ribosome biogenesis with RNAPol1 inhibitors was sufficient to redistribute MINA from the nucleolus to the nucleoplasm (Figures 3B and S4A)^9^. Publicly available SILAC proteomic data demonstrate that the response to RNAPol1 inhibition is rapid and dynamic (Figure 3C)^22^. Conversely, overexpression of the RNAPol1 activator RRN3 was sufficient to enrich some MINA in the nucleoli of confluent cells (Figure S4B). Nucleophosmin (NPM1) is a highly abundant nucleolar protein involved in ribosome biogenesis and a major MINA-binding partner^9, 13^. To test whether NPM1 contributes to nucleolar sequestration of MINA we targeted it with siRNA in subconfluent Caco-2 cells. Consistent with the hypothesis, loss of NPM1 expression was sufficient to redistribute MINA to the nucleoplasm (Figure S4C). Overall, our localization analyses suggest that nucleolar enrichment of MINA is dependent on elevated levels of ribosome biogenesis and its nucleolar interactome. In cells where these determinants are lost or suppressed, MINA is redistributed to the nucleoplasm, where (we hypothesize) it undertakes an alternative function in epithelial membrane biology.

**Figure 3.**
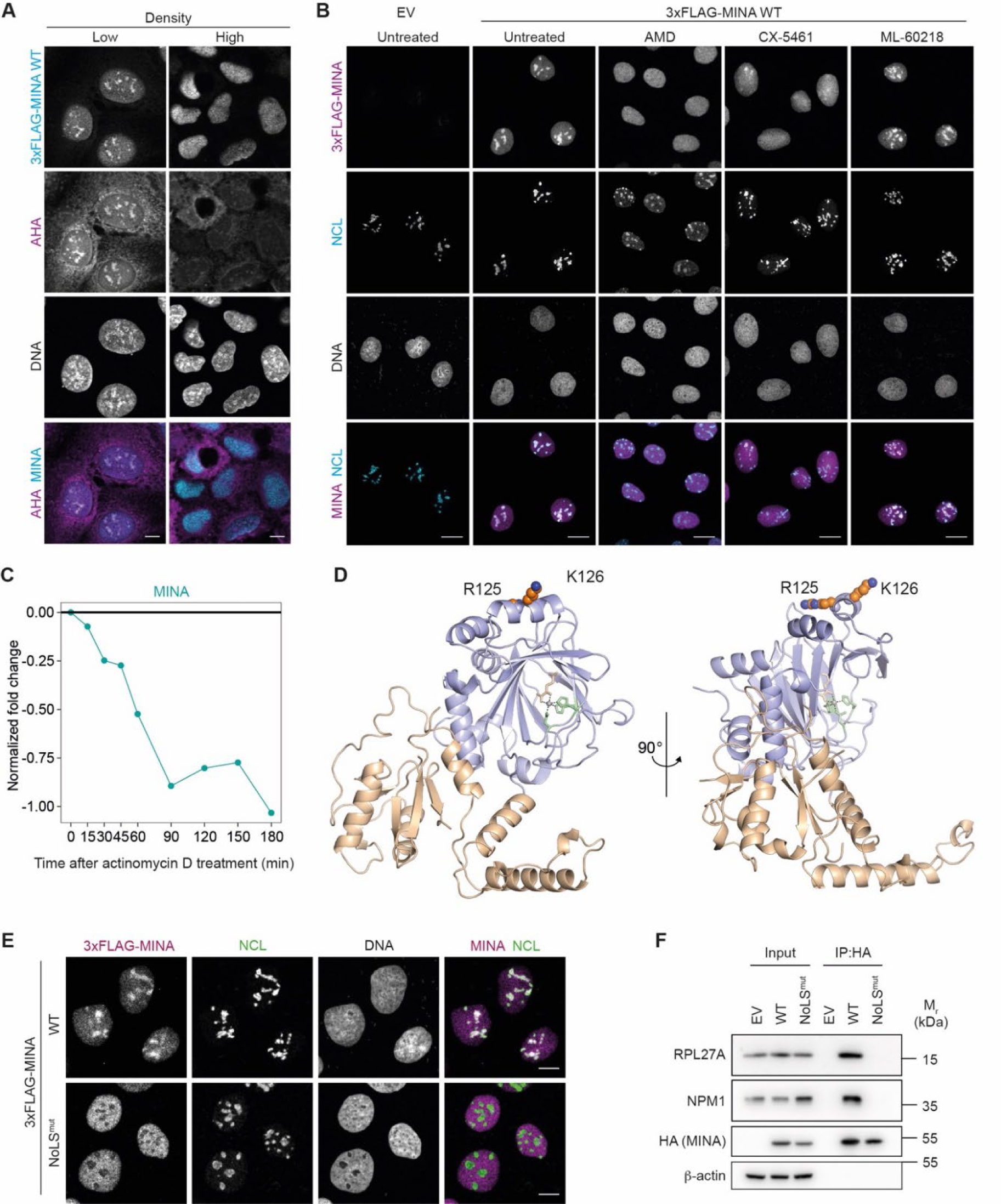
Determinants of subnuclear MINA localization. **(A)** 3xFLAG-MINA colocalizes with newly synthesized proteins in the nucleoli of subconfluent cells. Stable Caco-2 cells expressing wildtype (WT) 3xFLAG-MINA and seeded at a high or low density were treated with methionine analogue L-Azidohomoalanine (AHA, magenta) for 1 h before the cells were fixed and labelled using Click-IT chemistry. Cells were co-stained for MINA (cyan). Scale bars, 10 µm. **(B)** Nucleoplasmic translocation of MINA in response to RNA polymerase I inhibitors. Expression of 3xFLAG-MINA was induced in subconfluent Caco-2 cells using 0.1 µg/mL doxycycline for 48 h prior to treatment. Empty vector (EV) or WT 3xFLAG-MINA expressing cells were either untreated or treated with 40 ng/mL actinomycin D (AMD) or CX-5461 to inhibit RNA polymerase I for 4 h. Parallel samples were treated with RNA polymerase III inhibitor ML-60218 to determine the specificity of the observed effects. Cells were stained for MINA (magenta), nucleolin (NCL, cyan) and DNA (DAPI, gray). Scale bars, 20 µm. **(C)** SILAC proteomic data demonstrate that the nucleolar abundance of MINA declines rapidly in response to actinomycin D (see Anderson et al, main text ref ^22^). The authors treated HeLa cells with 1 µg/mL actinomycin D for the indicated times before mass spectrometry-based quantification of the nucleolar proteome. **(D)** Ribbon representation of a MINA monomer showing residues that contribute to MINA nucleolar localization (R125/K126) and the catalytic core in purple. Protein Data Bank (PDB) ID = 4BXF. Structures were visualized using PyMOL. **(E)** R125G/K126G mutations prevent MINA nucleolar localization. Caco-2 cells expressing either WT or 3xFLAG-MINA R125G/K126G (NoLS^mut^) were stained for MINA (magenta), nucleolar protein nucleolin (NCL, green) and DNA (DAPI, gray). Scale bars, 10 µm. Expression of 3xFLAG-MINA was induced by treatment with 0.1 µg/mL doxycycline for 48 h. **(F)** MINA NoLS^mut^ does not interact with nucleolar interactors RPL27A and NPM1. Anti-HA immunoprecipitates (IP) from extracts of HEK293T cells transiently expressing the indicated HA-MINA constructs were immunoblotted for RPL27A (17 kDa), NPM1 (33 kDa) and HA (MINA, 53 kDa). β-actin (42 kDa) was used as loading control. Input = cell extract prior to immunoprecipitation (for RPL27A and NPM1 the input is 1% of the IP, for HA and β-actin it is 50%). β-actin (42 kDa) was used as loading control. Uncropped western blot images are presented in source data.

We reasoned that a nucleolar localization mutant would support efforts to dissect nucleolar versus extra-nucleolar MINA functions. We searched the MINA primary sequence for nucleolar localization sequences (NoLSs)^23^ and identified two potential candidates, one at the N-terminus and a bipartite-like motif within residues 100-125 (Figure S5A). The latter was more highly conserved than the N-terminal sequence (Figure S5B). The basic residues at the C-terminal end of the conserved bipartite motif (R125/K126) are within an a-helix and their side chains are solvent-exposed, which are known determinants of NoLSs (Figure 3D)^24^. The C-terminal part of bipartite NoLSs have previously been shown to be involved in nucleolar targeting^24^. Therefore, we considered R125/K126 as residues of interest to experimentally test their contribution to MINA nucleolar localization. Consistent with predictions, an R125G/K126G mutant (NoLS) was unable to localize to the nucleolus (Figure 3E) and did not interact with the nucleolar targets RPL27A and NPM1 (Figure 3F). The R125G/K126G mutation retained nuclear localization (Figure 3E) and hydroxylase activity (Figure S5C), suggesting that MINA NoLS mutant is a viable tool for investigating the extra-nucleolar role of MINA.

### Extra-nucleolar MINA interacts with membrane palmitoylated protein 6 (MPP6)

To investigate how extra-nucleolar MINA regulates barrier function we next sought to identify extra-nucleolar MINA interactors. We hypothesized that a physiologically relevant extra-nucleolar MINA interactor might exist in multiple independent cell types and be enriched in NoLS versus wildtype MINA pulldowns. Therefore, we first immunopurified wildtype 3xFLAG-MINA from three diverse cell types (Caco-2, U2OS, and mouse embryonic fibroblasts) before trypsinolysis and MS-based protein identification. We removed contaminants present in control samples before applying strict filtering criteria to identify only the highest confidence and most abundant interactors. Cross-referencing the three datasets identified just four proteins in common, two of which were the known interactors RPL27A and NPM1. Next, we compared NoLS and wildtype MINA proteomic screens performed in Caco-2 cells to identify proteins with increased binding to extranucleolar MINA: We cross-referenced these candidates with the four proteins identified in multiple cell lines above, which identified one protein, MPP6 (Figure 4A). Because MPP6 is a member of the MAGUK superfamily of cell adhesion and polarity proteins (Figure S6)^25^, we considered it a viable candidate to mediate the extra-nucleolar functions of MINA in epithelial membrane biology. Therefore, we sought to study the MINA:MPP6 interaction in more detail.

**Figure 4.**
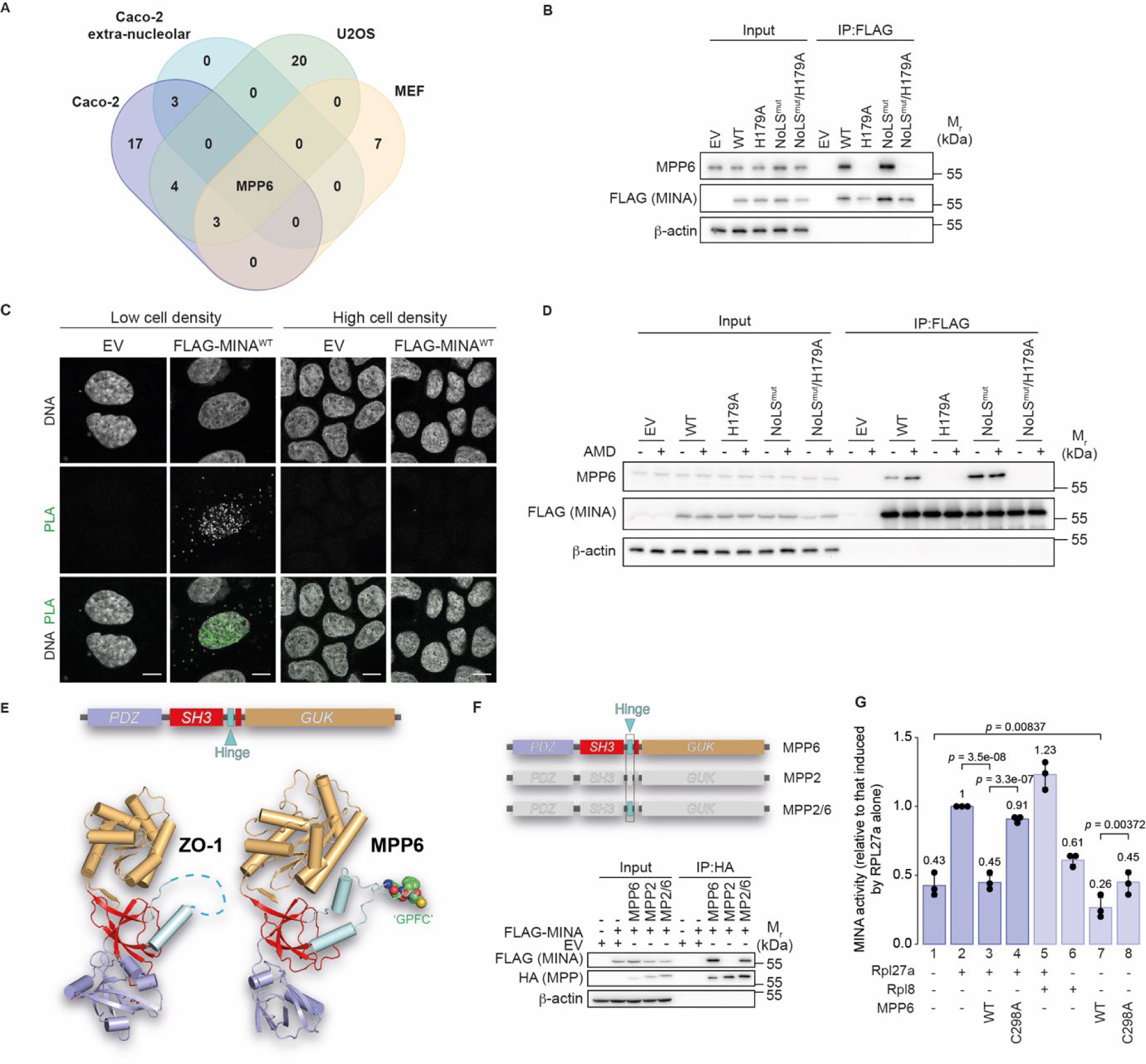
Extra-nucleolar MINA interacts with MPP6, a member of the ‘membrane-associated guanylate kinase’ (MAGUK) family. **(A)** Venn diagram showing the overlap between high stringency MINA-interactomes identified in Caco-2, U2OS and MEF cells. Note that the only common factor that was also upregulated with extra-nucleolar MINA (NoLS^mut^) was MPP6. **(B)** Validation of proteomic data. Anti-FLAG immunoprecipitates (IP) from extracts of stable Caco-2 cells expressing the indicated doxycycline-inducible 3xFLAG-MINA constructs were immunoblotted for endogenous MPP6 (61 kDa), FLAG (MINA, 53 kDa). For MPP6 the input is 0.6% of the IP, for FLAG and β-actin it is 31%. **(C)** Proximity ligation assays (PLA, green foci) indicate that MINA and MPP6 preferentially interact in the nuclei (gray) of subconfluent Caco-2 cells. Scale bars, 10 µm. **(D)** The interaction of MPP6 with MINA is suppressed by RNA polymerase I activity. Anti-FLAG IPs from extracts of stable SW620 cells expressing an empty vector (EV) or the indicated 3xFLAG-MINA constructs were immunoblotted for MPP6 (61 kDa) and FLAG (MINA, 53 kDa). Expression of 3xFLAG-MINA was induced with 0.2 µg/mL doxycycline for 24 h before the cells were treated with 40 ng/mL actinomycin D (AMD) for 4 h. Input = 1%. **(E)** MPP6 domain architecture. Top, Cartoon representation of the ‘PSG’ supramodule. Bottom, Comparison of the X-ray structure of the PDZ-SH3-GUK region from a canonical MAGUK protein ZO-1 (PDBID:3SHW) and the predicted structure of the equivalent region of MPP6 generated with AlphaFold. The GPFCG motif is highlighted in space-filling representation and is located at the tip of the unstructured ‘Hinge’ insert. The yellow ball represents residue C298. **(F)** The Hinge domain of MPP6 is sufficient for MINA binding. An MPP2 chimera that contains the Hinge domain of MPP6 (AAs 285-338) (‘MPP2/6’), but not wildtype MPP2, coprecipitates FLAG-MINA. Top, Graphical representation of the constructs used in the bottom panel. Bottom, Anti-HA immunoprecipitates from extracts of HEK293T cells expressing the indicated constructs were immunoblotted as indicated. For FLAG and β-actin the input is 10% of the IP, for HA it is 20%. **(G)** An MPP6 peptide containing the minimal determinants for MINA binding (SQFLEEKRKAFVRRDWDNS**GPFCG**TISSKKKKK) blocks RPL27A peptide (GRGNAGGL**H**HHRINFDKYHP)-induced MINA activity in vitro. An NO66 substrate peptide (Rpl8; NPVEHPFGGGNHQHIGKPSTI), which is known not to support MINA activity ^13^ was used as negative control. MINA activity was determined using an indirect coupled assay based on succinate production (see STAR Methods). Bars represent the average of n = 3 biological repeats. Averages are indicated above each respective bar. Error bars, SD. Statistical significance was determined using a one-way ANOVA and p values were calculated post-hoc using a pairwise t test. P values were adjusted using the Benjamini-Hochberg correction for multiple comparisons. Uncropped western blot images are presented in source data.

First, we independently validated the proteomic results in multiple cell types (Figures 4B, S7A and S7B). As expected, we observed a specific interaction between MPP6 and MINA that was increased in the nucleolar localization mutant. Intriguingly, the interaction was dependent on the catalytic activity of MINA, being completely abolished by an inactivating Fe(II)-binding mutation (H179A) (Figures 4B, S7A and S7B) (the role of MINA activity is investigated in more detail below). Next, we explored the specificity of the MINA:MPP6 interaction. Under conditions where endogenous MPP6 interacted with HA-MINA, it did not bind to the closely related hydroxylase NO66 (Figure S7C). Similarly, FLAG-MINA coprecipitates with MPP6-HA, but not with the highly related MPP2 (sequence identity = 65.2%) (Figure S7D). Endogenous MPP6 coprecipitates endogenous MINA (Figure S7E). Consistent with the MINA localization (Figure 1) and proteomic analyses (Figure 4), immunofluorescence and proximity ligation assays (PLA) indicate that 3xFLAG-MINA and MPP6-HA colocalize and interact in the nucleoplasm of subconfluent cells (Figures 4C and Figure S7F). Moreover, the PLA signal was reduced in confluent monolayers (Figure 4C), consistent with the translocation of MPP6 from the nuclear compartment to the membrane (Figure S7G). Unsurprisingly therefore, less MPP6 interacts with 3xFLAG-MINA in highly confluent monolayers of Caco-2 (Figure S7H). Increasing the pool of extra-nucleolar MINA in subconfluent cells (through RNApol1 inhibition, Figure 1) increased its interaction with MPP6 (Figure 4D). The NoLS mutant showed constitutively increased MPP6 binding, as expected, and was completely resistant to the effects of RNApol1 inhibition (Figure 4D). These data indicate that RNApol1 activity and ribosome biogenesis sequester MINA in the nucleolus of subconfluent cells, thereby suppressing its interaction with MPP6.

### The Hinge regions of MPP2 and MPP6 confer MINA binding specificity

As scaffold proteins, MAGUK proteins are characterized by the presence of several protein-protein interaction domains. These include a characteristic PDS-95/DLG/ZO-1 (PDZ) – Src homology 3 (SH3) – guanylate kinase (GUK) tandem motif (‘PSG’) (Figure S6) that can fold into a structural supramodule^26^. Folding requires a disordered ‘Hinge’ region that loops out of the non-canonical SH3 domain and that includes a polybasic sequence implicated in membrane localization and protein:protein interactions^27^. Modeling the tertiary structure of MPP6 using AlphaFold confirms the likelihood that the PDZ, SH3, and GUK domains of MPP6 form a PSG supramodule analogous to ZO-1 (Figure 4E). Furthermore, the AlphaFold model of MPP6 also predicts the presence of a Hinge insert within its non-canonical split SH3 domain.

To test the requirements of these MPP6 domains for MINA binding we analyzed a series of MPP6 truncation mutants in immunoprecipitation and immunofluorescence experiments (Figures S8A, S8B, and S8C). The results suggest that the minimal MPP6 domain required for MINA binding is within the non-canonical SH3 domain, i.e. amino acids 285-338, which contains the so-called Hinge domain (Figures 4E and S8B). To test whether this domain is sufficient for MINA binding we created an MPP2/6 chimera by swapping the Hinge domain of MPP2 (which does not bind MINA; Figure S7D) with the corresponding domain of MPP6 (Figure 4F, top). Immunoprecipitation analysis confirmed that the MPP6 Hinge domain is sufficient for MINA binding in this context (Figure 4F, bottom). To further narrow down the binding determinants, we next focused on the differences in primary sequence between MPP2 and MPP6 in this region. The sequence identity of this region is 67.9%, suggesting that the strikingly different MINA binding profiles are determined by the relatively few regions of low homology (Figure S8D). To test the importance of these regions we swapped residues within them in the context of the MPP2/6 chimera. A small insertion at the N-terminal end of the MPP2 Hinge domain was sufficient to prevent MINA binding (Figure S8E). Next, we alanine scanned residues 287-303 within the Hinge domain of full-length MPP6 and identified residues 295-299 (sequence GPFCG) as critically required for MINA binding (Figure S8F). These residues are situated at the predicted apex of the unstructured and solvent-exposed loop of the Hinge domain (Figure 4E), and are therefore likely to be available for protein:protein interactions. Together, these data are consistent with a site-specific interaction of MINA with a minimal sequence motif within the Hinge domain of MPP6.

### The Hinge region of MPP6 mediates a pseudo-substrate-like interaction with MINA

MINA interacts with MPP6 in an activity-dependent manner (Figures 4B, 4D, S7A and S7B). The interaction of MINA with the MPP6 Hinge domain retains its dependence on MINA hydroxylase activity (Figures S8G and S8H). Activity-dependence of binding can be an indicator that a hydroxylase interactor is a novel substrate^28, 29^. Therefore, we next explored whether MPP6 is hydroxylated by MINA.

MPP6-3xFLAG was overexpressed in the presence or absence of HA-MINA before immunoprecipitation (Figure S9A), in-solution chymotrypsin digest, and LC-MS/MS analysis. Near complete (98%) peptide-level sequence coverage of MPP6 was achieved in database searches that also considered a wide-range of residue-specific oxidations (Figure S9B). Consistent with the lack of homology between the known substrate RPL27A and the Hinge domain of MPP6 (Figure S9C) we did not detect histidyl hydroxylation in this region. Oxidized peptides encompassing the Hinge domain were detected that could be confidently assigned to W291 (Figures S9D and S9E). However, the stoichiometry was low and did not respond to MINA overexpression (Figure S9F). Neither subconfluent culture (where MINA binding is predicted to increase), or proteasomal inhibition, led to elevated W291 oxidation, even in the presence of MINA overexpression (Figure S9F). We note that tryptophan can undergo spontaneous chemical oxidation as an artefact of sample preparation for MS^30–32^.

In order to take an orthogonal approach that avoids the limitations of mass spectrometry analysis and artefactual oxidation we next used a coupled in vitro enzyme assay. Although the currently assigned hydroxylase activity of MINA is restricted to histidyl residues, we noted the absence of a histidine in the minimal domain within MPP6 that is required for MINA binding (residues 295-299). Since some closely related JmjC-hydroxylases can target multiple amino acid types^33–36^, we first explored the potential for wider biochemical specificity of MINA, including the amino acids in this region of MPP6 (Gly, Pro, Phe, and Cys). To do this we used RPL27A peptides with the corresponding amino acid substitutions at the H39 target residue: Unlike the wildtype RPL27A peptide sequence, neither H39G, H39P, H38F or H39C peptides were able to support MINA activity (Figure S10A). Similar data were obtained using H39W-, H39S-, H39N-, H39D-, H39K-, and H39R-substituted peptides. These data indicate that MINA does not have a broad biochemical activity and is unlikely to hydroxylate residues found within the MPP6 minimal binding domain. To explore this directly, we tested three MPP6 peptides of varying length and position that overlap the GPFCG motif and wider Hinge region of MPP6 (which is sufficient for activity-dependent MINA binding, Figure 4F) (Figure S10B). Unlike the RPL27A peptide, none of the MPP6 peptides supported MINA activity (Figure S10C).

Overall, therefore, our cell-based and in vitro studies did not identify a MINA-dependent hydroxylation of MPP6. We cannot rule out the presence of a modification that was below the limits of detection, the catalysis of a previously undescribed oxidative modification, or subsequent processing of an oxidized species. However, it is also possible that, unlike the activity-dependent interactions of related JmjC hydroxylases that led to novel substrate assignments^28, 29^, the activity-dependent interaction described here is unique. For example, the interaction of MPP6 with MINA may require the integrity and availability of the catalytic pocket, rather than being ‘activity-dependent’ per se. In line with MPP6 binding directly to the MINA catalytic pocket, an MPP6 peptide containing the GPFCG motif prevented RPL27A-induced MINA activity (Figure 4G; compare 2^nd^ and 3^rd^ bar) and significantly reduced its ‘uncoupled’ turnover (compare 1^st^ and 7^th^ bar). Both effects were specific as they were not observed with a control peptide (NO66 substrate Rpl8) (compare 2^nd^, 3^rd^, and 5^th^ bars, and 1^st^, 6^th^ and 7^th^ bars). Furthermore, both effects of the MPP6 peptide were ablated by the C298A substitution (compare 3^rd^ and 4^th^ bar, and 7^th^ and 8^th^ bar) (Figure 4G), which is sufficient to block binding of MPP6 to MINA (Figure S8F). Overall, our data are consistent with the possibility that the solvent exposed C298 residue of the MPP6 Hinge domain binds to the MINA catalytic pocket in a pseudo-substrate-like manner.

### MINA regulates MPP6 expression and localization

Although we did not identify a modification arising from MINA binding to MPP6, we were intrigued that the function of the interaction might be related to the role of the Hinge domain in other MAGUKs, i.e. membrane localization and protein stability^27, 37^. MINA knockdown caused a reduction in membrane staining and increase in nuclear localization of MPP6 in low-density cultures (Figures 5A and 5B), in parallel with a reduction in MPP6 protein expression (Figure 5C). Consistent with these changes being a direct consequence of the interaction with MINA, a constitutively expressed MPP6 mutant that fails to bind MINA (C298A) was nuclear localized and underexpressed (Figures 5D and 5E). These data suggest that the interaction of MINA with MPP6 in the nucleus (Figures 4C) promotes the subsequent localization of MPP6 to the plasma membrane.

**Figure 5.**
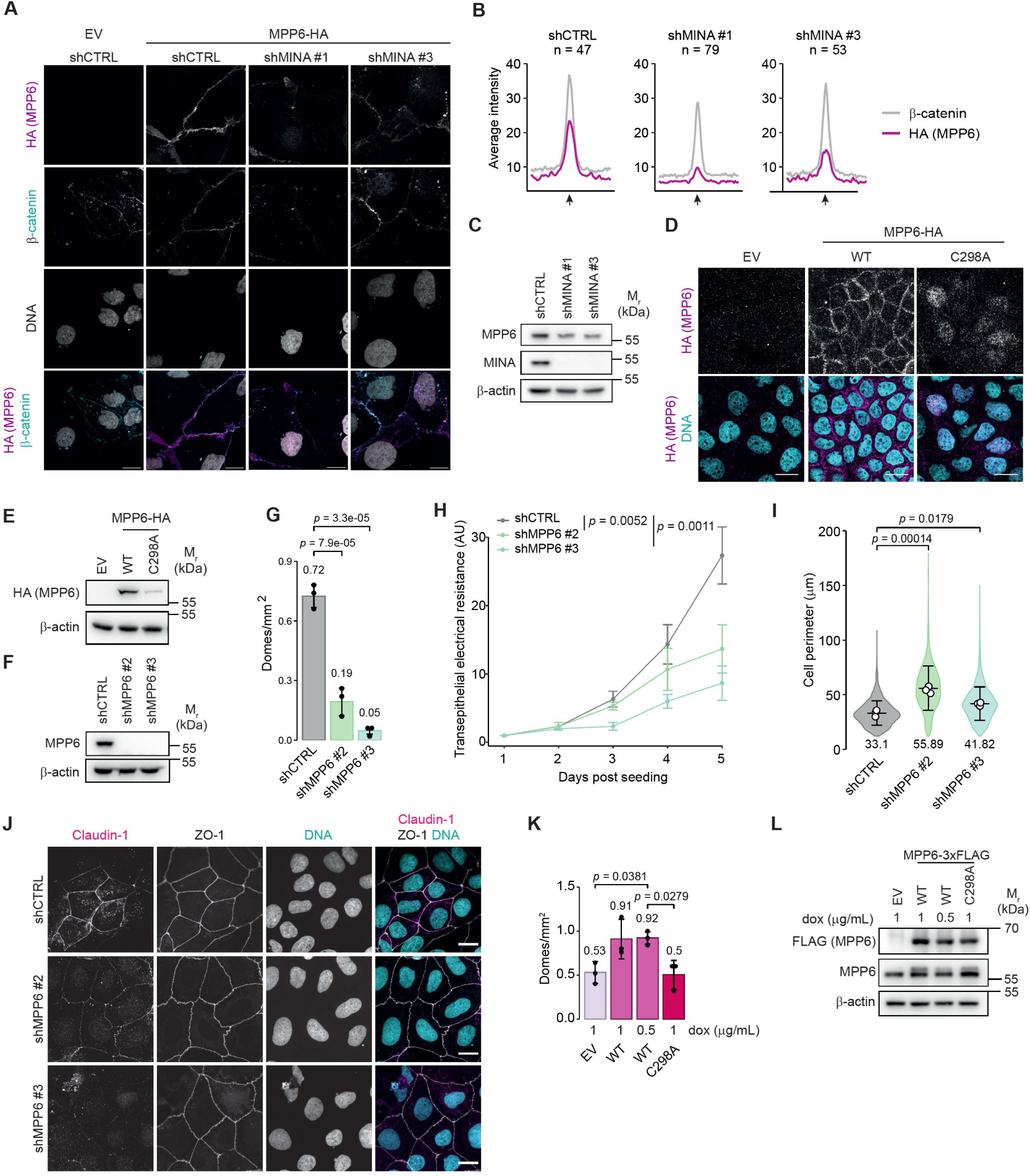
MPP6 promotes tight junction integrity and epithelial barrier function. **(A)** MINA depletion reduces membrane localization and increases nuclear distribution of MPP6. Representative immunofluorescence images of subconfluent Caco-2 cells expressing the indicated shRNAs and stained for HA (magenta), β-catenin (cyan) to mark the membranes and DNA (DAPI, gray). Scale bars, 20 µm. **(B)** Reduced membrane staining of MPP6-HA in MINA-depleted cells under conditions presented in (A). Line graphs indicate the average fluorescence intensity of a line drawn across the membrane connecting two cells, measured with Fiji. The average of n = 3 biological experiments is shown. The total number of images analyzed for each condition is shown above each respective graph. β-catenin was used as membrane marker. **(C)** MINA depletion reduced MPP6 expression in subconfluent Caco-2 cultures. Whole cell extracts from cells expressing the indicated shRNAs were immunoblotted for MPP6 (61 kDa) and MINA (53 kDa). **(D)** Altered localization of HA-tagged MINA-binding mutant MPP6^C298A^. Stable Caco-2 cell lines constitutively expressing an empty vector (EV), wildtype (WT) or MPP6-HA C298A were stained for HA (MPP6, cyan) and DNA (DAPI, cyan). Scale bars, 20 µm. **(E)** Reduced expression of HA-tagged MINA-binding mutant MPP6^C298A^. Whole cell extracts were immunoblotted for HA (MPP6, 61 kDa). **(F)** Representative immunoblot from whole cell extracts of stable Caco-2 cells expressing the indicated shRNAs showing efficient depletion of MPP6 (61 kDa). **(G)** MPP6 depletion reduces dome formation. Bar graph showing the average number of domes per mm^2^ of n = 3 independent experiments. The average of each condition is shown above each respective bar. **(H)** Delayed establishment of transepithelial electrical resistance in MPP6-depleted cells. Line graph depicts the average of three technical repeats of one biological repeat. The experiment was repeated 4 times with similar results. Statistical significance was determined on day 5. **(I)** Increased perimeter of cells in MPP6-depleted monolayers. Violin plot represents the data from n = 3 experiments. Average of each condition is shown underneath each respective violin. n = 2612 (shCTRL), n = 1450 (shMPP6 #2) and n = 1958 (shMPP6 #3) cells were quantified. **(J)** Altered localization of Claudin-1 following MPP6 knockdown. Representative images of Caco-2 monolayers expressing the indicated shRNAs were stained for Claudin-1 (magenta), ZO-1 (gray) and DNA (DAPI; cyan). Scale bars, 20 µm. **(K)** MPP6-induced dome formation is dependent on C298. Expression of wildtype (WT) or MINA-binding mutant C298A MPP6-3xFLAG were induced with doxycycline (dox) at the indicated concentrations. Note that a reduced concentration of dox (0.5 µg/mL) was included for WT to allow fair comparison to the reduced expression levels observed for C298A (as also presented in (E)). Domes were counted 4 days post seeding. **(L)** Cell extracts of samples treated as in (K) were immunoblotted for FLAG (MPP6) and MPP6 (61 kDa). (G-I and K) Statistical significance was determined using a one-way ANOVA and p values were calculated post-hoc using a Dunnett’s multiple comparisons test. Error bars, SD. (C, E, F and L) β-actin (42 kDa) was used as loading control. Uncropped western blot images are presented in source data.

### MPP6 is required for the timely establishment of cell-cell junctions

Considering the effect of MINA knockdown on barrier function (Figure 1) and MPP6 membrane localization and expression (Figures 5A and 5B), we hypothesized that MPP6 knockdown would phenocopy MINA loss of function. Indeed, shRNA knockdown of MPP6 severely impaired dome formation and TEER (Figures 5F, 5G and 5H) and increased the length of the cell perimeter (Figures 5I), with concomitant reduction in membrane staining and expression of Claudin-1 (Figures 5J, S11A and S11B). Similar to MINA knockdown, this occurred in the absence of gross changes in cell proliferation (Figure S11C). To explore the effect of the MPP6 C298A mutant we switched to a tunable doxycycline-inducible system, due to the known confounding effects of overexpressing membrane/polarity proteins in some circumstances, and the differential expression of constitutively expressed wildtype and C298A MPP6 (Figure 5E). Using this system, we observed that modest expression of MPP6-3xFLAG was sufficient to promote dome formation (Figures 5K and 5L). The MPP6 C298A mutant, which impairs MINA binding, was completely inactive in this regard (Figure 5K and 5L).

### The MINA-MPP6 pathway is required for normal 3D growth and morphogenesis

To investigate the importance of the MINA-MPP6 pathway in 3D growth and tissue morphogenesis, Caco-2 cells expressing doxycycline-induced control, MINA or MPP6 shRNA were seeded as single cells in Matrigel and grown for 7 days to form polarized spheroids with a fluid-filled lumen^17^. Depletion of MINA or MPP6 was sufficient to interfere with spheroid morphology including lumenogenesis (Figures 6A,B and S12A,B), size (Figure 6C,D), and growth (Figure 6E,F). Immunofluorescence analysis revealed a diffuse staining pattern of Claudin-1 and ZO-1 and altered nuclear morphology (Figure 6G,H). Although these observations may be consistent with some level of defective polarity, a luminal surface was still largely maintained in these spheroids (Figure 6G,H). Consistent with the 2D data presented above, MINA and MPP6 depletion also reduced Claudin-1 levels (Figure S12C,D). Expression of the MINA binding defective MPP6 mutant (C298A) phenocopied MINA and MPP6 loss of function, causing reduced lumenogenesis, decreased spheroid size, and diffuse localization of ZO-1 (Figure 6I).

**Figure 6.**
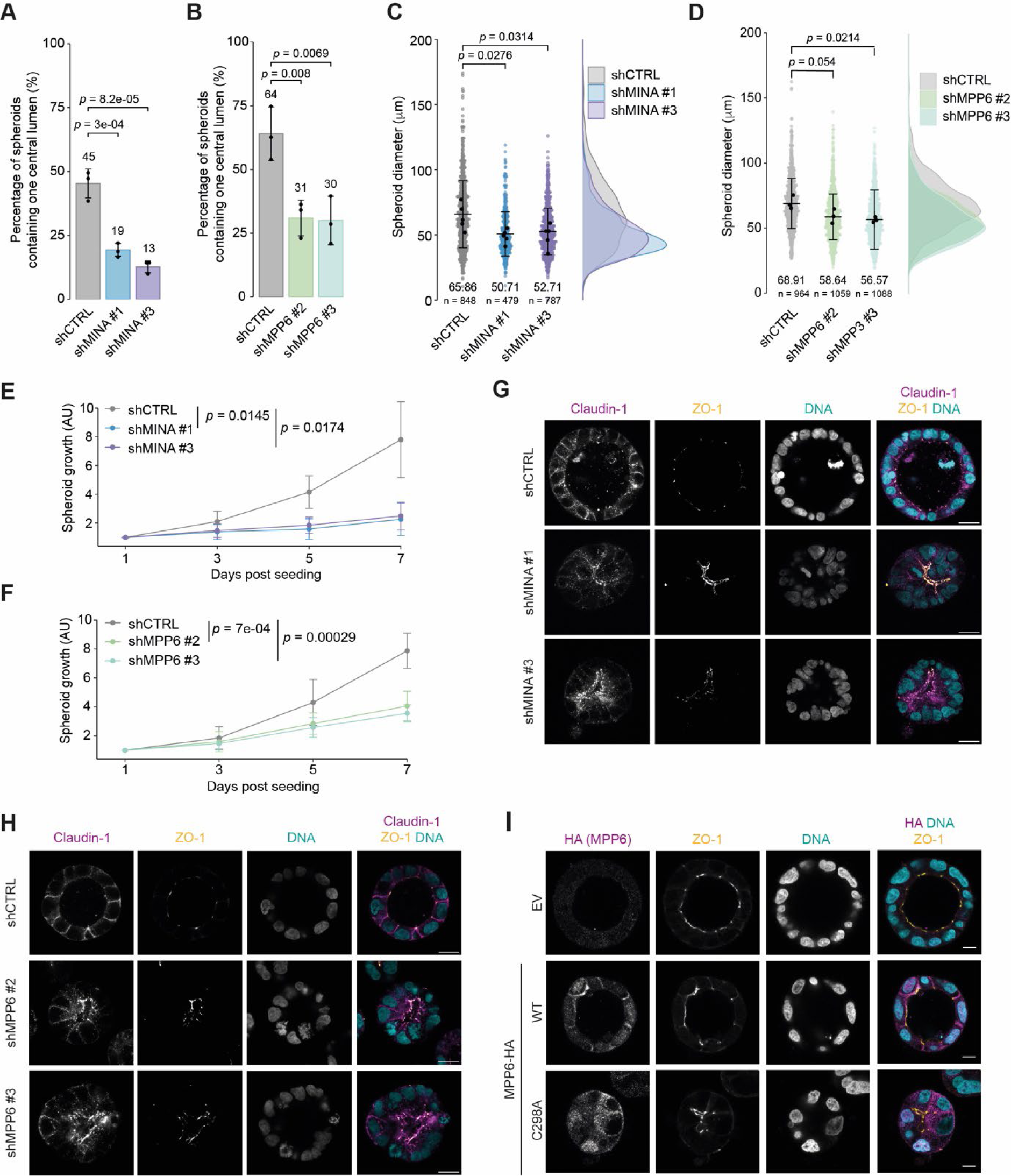
The MINA-MPP6 pathway is required for spheroid lumenogenesis, growth and polarity. **(A and B)** Quantification of lumen formation in spheroids grown from stable Caco-2 cells expressing the indicated MINA (A) or MPP6 (B) shRNA sequences. Bars represent the average percentage of spheroids with one central lumen of n = 5 (C) and n = 3 (D) independent experiments. The average of each condition is shown above each respective bar. **(C and D)** Quantification of size of spheroids grown from stable Caco-2 cells expressing the indicated MINA (C) or MPP6 (D) shRNA sequences. Horizontal lines represent the average spheroid diameter from n = 3 independent experiments. The average of each condition and the number of spheroids counted are shown below each respective beeswarm. **(E and F)** MINA (E) and MPP6 (F) depletion reduces spheroid growth. Growth of spheroids was analyzed using a CellTiter-Glo Luminescent Cell Viability Assay kit after 3, 5 and 7 days in culture. Presented values are the average of n = 4 independent experiments. AU, arbitrary unit. **(G and H)** MINA (G) and MPP6 (H) depletion prevent appropriate membrane localization of Claudin-1 and ZO-1. Representative immunofluorescence images of 3D Caco-2 spheroids expressing the indicated shRNAs and stained for Claudin-1 (magenta), ZO-1 (yellow), and DNA (DAPI, cyan). Scale bars, 20 µm. **(I)** MPP6 C298A overexpression phenocopies MINA and MPP6 knockdown. Single cells were grown to spheroids for 7 days, fixed and stained for HA (MPP6, magenta) and tight junction marker ZO-1 (yellow). Nuclei were visualized with DAPI (cyan). Scale bars represent 10 μm. **(A – D)** Statistical significance was determined using a one-way ANOVA and p values were calculated post-hoc using a Dunnett’s multiple comparisons test. Error bars, SD.

Overall, our data are consistent with MPP6 acting in a pathway downstream of extra-nucleolar MINA that regulates cell adhesion, barrier function, and morphogenesis. Because this pathway is suppressed by RNApol1 activity and the nucleolar proteome, the work identifies a novel link between ribosome biogenesis and epithelial membrane biology.

## Discussion

Here in this paper, we have uncovered the extra-nucleolar function of the protein hydroxylase MINA, demonstrating, for the first time, that it controls epithelial TJ integrity, barrier function, and morphogenesis through its interaction with the MAGUK protein MPP6. In doing so we have discovered that cellular growth status and ribosome biogenesis are functionally coupled to the epithelial plasma membrane, via a novel nuclear partnership between MPP6 and the 2OG-oxygenase MINA. The work raises the intriguing possibility that ribosome biogenesis might also couple to membrane biology in other ways, and that other growth responsive cellular processes might sink proteins that would otherwise suppress such roles. Could the uniquely nutrient-dependent 2OG-oxygenase family play a more general role in signaling between subcellular compartments to couple cell systems controlling cell growth and differentiation?

Although 2OG-oxygenases are still considered an emerging enzyme family, relative to others (such as kinases), they have already been implicated in a variety of essential cellular systems, from oxygen sensing and epigenetics to protein translation and collagen synthesis^38, 39^. These enzymes have not previously been demonstrated to play a role in regulating cell polarity and adhesion through the direct interaction with peripheral membrane proteins, however. Although the interaction with MPP6 was entirely dependent on the integrity of a critical Fe(II)-binding residue within the MINA catalytic pocket (Figure 4), it did not result in a detectable hydroxylation event (Figures S9 and S10). Although more work is required to definitively conclude that a modification is not catalyzed, it is also possible that MPP6 is sensing the availability of the catalytic pocket as an indicator of the absence of RPL27A and/or co-factors. Indeed, computational investigation of potential MPP6 interaction modes suggests that the GPFCG region at the tip of the Hinge domain (Figures 4E and S8F) might directly bind the MINA active site through formation of β-turn like structures (Figure S13). Although ‘pseudo-substrates’ have been identified for other enzyme classes, this may be one of the first potential candidates that we know of for the 2OG-oxygenase family. As such, the work is relevant to the wider field: Protein hydroxylases, including Hypoxia Inducible Factor prolyl hydroxylases, may be at risk of substrate misassignment^31^, in part because of the artefactual oxidation of residues such as tryptophan during MS sample preparation, as also demonstrated here (Figure S9). Our work raises the possibility that some proteins that interact with the catalytic domain of 2OG-oxygenases may be doing so in a pseudo-substrate-like manner, rather than as prime substrates.

Through our phenotypic and proteomic analyses of MINA, we have helped clarify the function of the poorly characterized human MAGUK protein MPP6. The *drosophila* homologue of MPP6 (*Varicose*) has been more extensively studied than its human counterpart: *Varicose* is essential for the assembly and function of septate junctions (which are thought to be the equivalent of vertebrate TJs) and for normal tracheal development^40, 41^. A similar role for MPP6 in TJ function in human cells is supported by our results showing that its loss of function causes reduced dome and TEER formation and decreased Claudin-1 expression (Figure 5). Under conditions of MINA or MPP6 knockdown we did not observe widespread mis-localization of tight or adherens junction markers however, particularly in 2D. Similar observations were made with *Varicose*^40^ and with other human polarity proteins, including MPP7, Scribble and Par3^42–44^. The prevailing hypothesis is that such proteins are not primarily involved in the establishment of apico-basal cell polarity, but rather in ensuring and maintaining junctional functionality. MINA appears to promote this function at least in part by facilitating MPP6 membrane localization (Figures 5A and 5B), which could in turn support normal expression (Figure 5C), as observed for other peripheral membrane proteins^32^. How MINA achieves this is not yet clear, although it seems likely that this is a consequence of the direct interaction with the Hinge domain of MPP6 in the nucleus. Although the function and interacting partners of the MPP6 Hinge domain are poorly characterized, mutagenesis screens in the fly model demonstrate it is essential^40^. The Hinge domain of related human MAGUK proteins have been implicated in membrane localization, protein-protein interactions, and proteasomal degradation^27, 37^, consistent with the results described here. Further work is required to explore how extra-nucleolar MINA supports the translocation of nuclear MPP6 to the plasma membrane to function in TJ biology.

TJs are critical for normal physiology and have been heavily implicated in disease^45–47^. Their barrier function plays a critical role in regulating permeability to small molecules, immune cells, pathogens, and parasites, and as such has been implicated in a wide variety of diseases across multiple tissues, including colitis, infection, edema, and allergy^45–47^. In light of the results presented here, it will be of interest to study the role of the MINA-MPP6 pathway in these contexts. A single-nucleotide polymorphism in the MINA gene is associated with increased risk of asthma in children^48^, and MINA knockout reduces the allergic response to inhaled allergens in mice^49^. MINA knockout mice also show increased clearance of parasitic gastrointestinal nematode worms^50^, which in some contexts can be associated with increased barrier permeability that supports the ‘washer/sweeper’ effect and/or immunological access^51, 52^. Whether a deregulated MPP6-TJ pathway could play a role in these contexts is of interest and warrants future investigation.

Alterations in cell:cell junctions and adhesion is a hallmark of cancer^53^. Loss of tumor suppressor genes in TJ complexes and signaling pathways disrupts their fence function, resulting in abnormal polarity and reduced cell:cell adhesion, which together disrupt morphogenesis and promote migration/invasion^54^. Although several mechanisms have been described by which TJs are rendered dysfunctional in epithelial cancers, it is likely that others remain to be discovered: We propose that the MINA-MPP6 pathway is one such candidate that is worthy of further investigation, for the following reasons. Firstly, although MINA has been proposed as a biomarker and drug target in a range of tumor types (reviewed in^55^), there is emerging evidence supporting a paradoxical and poorly characterized role in tumor suppression^15, 16^. Secondly, low MPP6 expression is associated with poor prognosis in some cancers, and the MPP6 gene is located in a chromosomal region (7p15-21) altered in several tumor types^56–63^. Thirdly, MPP6 interacts with Cell Adhesion Molecule 1 (CADM1), an immunoglobulin superfamily transmembrane protein that is a tumor suppressor in a wide variety of cancers^64^. Importantly, CADM1 loss-of-function is associated with reduced Claudin-1 expression and barrier function^65, 66^, which phenocopies MINA and MPP6 knockdown (Figures 1 and 5). Future work should focus on the mechanisms by which the MINA-MPP6 pathway may be inactivated in cancer and the consequences of this on the barrier and fence functions of TJs. Furthermore, the work presented here suggests that an important mechanism by which the MINA-MPP6 pathway could be suppressed is through oncogene-induced nucleolar sequestration of MINA due to elevated RNAPol1 activity and ribosome biogenesis. RNAPol1 inhibitors are being considered as novel chemotherapeutics (reviewed in^67^): Our hypothesis would predict that such treatments might not only negatively impact growth and viability, but also help to ‘normalize’ cell:cell adhesion and polarity in some contexts.

## Methods

### Plasmid constructs and cloning procedures

Coding sequences of MINA, MPP6 and NO66 were PCR cloned from plasmid DNA or cDNA into pFLAG (Sigma), pEF6 (Invitrogen), pTIPZ, pIPZ or pIHZ vectors. pIPZ and pTIPZ are derivatives of the pGIPZ and pTRIPZ lentiviral vectors (Open Biosystems) that lack green fluorescent protein (GFP) and red fluorescent protein (RFP), respectively. The coding sequences of MPP2 and MPP2/6 hybrids containing C-terminal HA-tags and restriction sites required for cloning into the pIPZ vector were obtained from Thermo Scientific as GeneArt String Fragment. Non-targeting control (ULTRA-NT#4), MINA (ULTRA-3406274, ULTRA-3406273) and MPP6 (ULTRA-3337606, ULTRA-3337607) shRNA sequences in pZIP-TRE3G vectors were obtained from TransOMIC. PCR reactions were driven by the Phusion High Fidelity DNA polymerase (New England Biolabs (NEB), M0530). Primers were designed to contain a C- or N-terminal tag and the required restriction sites for subsequent ligations. Constructs were confirmed by sequencing, carried out by Source Bioscience. For nucleotide exchanges, site-directed mutagenesis was performed using a Phusion Site-directed Mutagenesis kit (Thermo, F541). 5’ termini of the primers were phosphorylated using T4 polynucleotide kinase (PNK, Thermo, EK0031) prior to the reactions. All constructs were confirmed by sequencing.

### Cell culture

HEK293T, Caco-2, SW620 and U2OS cells were cultured in Dulbecco’s Modified Eagle Medium (DMEM, Gibco) supplemented with 10% (v/v) fetal bovine serum (FBS, Sigma) and 100 international units/mL (IU/mL) penicillin and 100 µg/mL streptomycin (Gibco). Cell culture medium for stable cell lines was supplemented with 1 µg/mL puromycin (Gibco). For 3D spheroid cultures, Caco-2 cells were seeded into Matrigel-coated (Corning, 35621) chamber slides and the culturing medium was supplemented with 4% (v/v) Matrigel. All cell lines were grown at 37°C in a humidified atmosphere with 5% (v/v) CO_2_. All cell lines were regularly tested for mycoplasma contamination using a LookOut® Mycoplasma PCR Detection kit (Sigma, MP0036).

Where doxycycline-inducible shRNA cell lines were used for experiments, shRNA expression was induced using 1 µg/mL doxycycline (Sigma, D9891) for 72 h before seeding. Subsequently, the culturing medium was refreshed, and fresh doxycycline was added every 48-72 h.

### Transient transfection

Plasmid DNA transfections were performed using FuGENE^®^ 6 transfection reagent (Promega). Briefly, the transfections mix was made up in Opti-MEM™ (Gibco, 31985070) using a ratio of DNA to transfection reagent of 1:3 and incubated for 15-30 min at room temperature (RT) before the complexes were added to the cells. 500 ng of DNA was used per mL of culturing medium. Cells were seeded 24 h before transfection and lysed or fixed 48 h after transfection.

### Generation of stable cell lines

Stable expression cell lines were generated by transduction with lentivirus. For lentivirus production, HEK293T cells were transfected with the plasmid of interest (50%), the packaging vector psPAX2 (35%) and envelope vector pMD2.G (15%) as described above. The lentivirus-containing supernatant was collected 48 h after transfection, passed through a 0.45 µM polyethersulfone (PES) filter, diluted 1:1 with fresh medium and added to the recipient cells for 48 h. Thereafter, transduced cells were selected using puromycin (2 µg/mL final concentration, Gibco).

### Metabolic labelling of newly synthesized proteins

Protein synthesis was monitored with L-azidohomoalanine (AHA) using a Click-iT™ Nascent Protein Synthesis kit (Thermo Scientific) according to the instructions of the manufacturer. Briefly, cells were incubated with methionine-free medium for 1 h before AHA was added at a final concentration of 50 mM. After 1 h, cells were fixed and permeabilized as described below. Next, Alexa Fluor 595 Alkyne diluted in Click-iT^®^ Cell Reaction Buffer (Thermo Scientific) at a final concentration of 2.5 mM was conjugated to any incorporated AHA.

### Proximity ligation assay

Duolink® proximity ligation assay kits were purchased from Sigma-Aldrich and performed according to the manufacturer’s instructions.

### Immunofluorescence staining

For immunofluorescence studies of 2D cultures, cells were grown on poly-D-lysine-coated or uncoated glass coverslips and fixed or 4% (w/v) paraformaldehyde (PFA) for 15 min at RT with methanol for 10 min at -20°C. Where methanol was used, it is stated in the figure caption. Cells fixed with PFA were subsequently permeabilized using 0.1% (v/v) Triton in PBS for 10 min at RT. Next, cells were blocked with 1% (w/v) bovine serum albumin (BSA) in PBS for 1 h at RT and subsequently incubated overnight (ON) at 4°C with primary antibodies diluted in 1% (w/v) BSA in PBS. The following primary antibodies were used: FLAG (Sigma, F1804, 1:1,000), HA (Covance, MMS-101P, 1:500), HA (Cell Signaling Technology (CST), C29F4, 1:500), nucleolin (Santa Cruz Biotechnology (SCB), sc-13056, 1:200-1:1,000), nucleophosmin-1 (SCB, sc-6013-R, 1:200), MINA (Invitrogen, 39-7300, 1:100), Claudin-1 (Abcam, ab211737, 1:500), β-catenin (BD Transduction Laboratories, 610153, 1:500). The next day, coverslips were washed with 1% (w/v) BSA in PBS and incubated with secondary antibodies diluted in 1% (w/v) BSA in PBS for 1 h at RT. Alexa Fluor 488, 555 or 633 secondary antibodies were purchased from Invitrogen and used at a final concentration of 1:1,000. Nuclei were stained by incubation with 0.5 – 1 µg/mL 4’,6-diamidino-2-phenylindole (DAPI, D1306, Invitrogen) in PBS for 10 min at RT, and the coverslips were mounted with Prolong Gold antifade mountant (Molecular Probes) onto microscope slides.

Caco-2 cells grown as 3D cultures were fixed with 4% (w/v) PFA/5% (w/v) sucrose in PBS for 20 min at RT, washed 3 times with PBS and incubated with 100 mM glycine in PBS for 20 min at RT to quench unreacted aldehydes. Subsequent permeabilization, blocking and staining were performed as described above. Stained spheroids were stored in PBS at 4°C until they were imaged.

### Image acquisition, processing, and quantification

Confocal images were acquired using a Zeiss LSM 780 or 880 laser scanning confocal system equipped with a 20x, 40x or 60 C-Apochromat 1.2 W Korr M27 objective. Immunofluorescence images of similarly stained experiments were acquired with identical laser power and detector gain. Images were processed using Fiji software (https://fiji.sc) and arranged in Adobe Illustrator. Brightness and contrast were adjusted using only linear operations applied to the entire image.

Cell perimeters were measured using the CellMorph plugin for Fiji (https://github.com/viboud12/CellMorph). Ten random images were taken for each condition in each experiment. Quantification was based on ZO-1 staining.

Phase-contrast images were acquired using an EVOS™ FL color imaging system equipped with a 4x or 10x Plan PH2 objective. To quantify domes, a total of 30 phase-contrast images were taken per condition per experiment, 4 or 5 days after seeding when the cells had formed a confluent monolayer. To quantify spheroids, a total of 10 images were taken per condition per experiment, 7 days after seeding.

### Multiplex Immunofluorescence Staining

4 µm formalin fixed paraffin embedded (FFPE) normal human colon were sectioned on to TOMO hydrophilic adhesive microscope slides (Matsunami).

Following antibody validation, fully automated multiplex immunofluorescent staining was performed on the Ventana Discovery Ultra platform (Roche Tissue Diagnostics, RUO Discovery Universal V21.00.0019). Primary-secondary-antibody pairs were as follows: Ki67 (Roche Tissue Diagnostics, 05278384001), OmniMap-anti-Rb HRP (Roche Tissue Diagnostics); pan-CKAE1-AE3 (Leica Biosystems, NCL-L-AE1/3,1:250), OmniMap-anti-Ms HRP (Roche Tissue Diagnostics); MINA (Thermo Fisher, 409500, 1:50), OmniMap-anti-Ms HRP (Roche Tissue Diagnostics); UBF1(Abcam, Ab75781, 1:50), UltraMap-anti-Rb HRP (Roche Tissue Diagnostics). Fluorescent detection was performed using an Opal fluorophore tyramide-based signal amplification system (Akoya Biosystems) with the following combinations. Ki67: Opal 570 (1:200); pan-CKAE1-AE3:Opal 620 (1:100) : MINA:Opal 690 (1:200); UBF1:Opal 520 (1:200).

A nuclear Hoechst 33342 counterstain (Sigma, 14533, 1:50) was applied to each sample for nuclear detection.

### Multiplex Immunofluorescence Image acquisition and Analysis

Multispectral whole slide images were acquired on a Vectra Polaris imaging system (Akoya Biosystems) using a 40x objective. Spectral unmixing was performed using a spectral library generated from single fluorescence slides of each protein marker with their assigned fluorophore dye, a DAPI only counterstain slide, and an unstained normal colon tissue section. The resulting images were analyzed using Visiopharm software (version 2022.12.0.12865). A bespoke deep learning algorithm was trained using DAPI, UBF1, and Ki67 features to detect nuclei and nucleoli in the epithelial compartment. Mean pixel intensities for each marker, in addition to X,Y coordinates, were exported for subsequent statistical analysis.

Single cell measures obtained from Visiopharm were analysed using R (version 4.2.2) with tidyverse packages (version 2.0.0). Per crypt binned quartile cellular distances were determined by calculating their perpendicular point along the crypt axis. Nucleolus:Nuclear ratio calculated as mean nucleolar per pixel intensity divided by mean nucleoplasm per pixel intensity.

### Transepithelial electrical resistance measurements

Caco-2 cells were seeded onto cellQART® permeable support inserts (0.4 mm pore size). The transepithelial electrical resistance (TEER) was measured using an EVOM2 epithelial voltohmmeter (World Precision Instruments, Inc.) and a STX2 electrode (World Precision Instruments, Inc.) according to the manufacturer’s instructions. Background resistance was measured using cell-free inserts. Three measurements were taken for each sample at each time point.

### Whole cell extracts

Cells were washed and harvested in ice-cold PBS, pelleted by centrifugation at 234 × g and lysed by rotation at 4°C for 1 h in JIES buffer (100 mM NaCl, 20 mM Tris-HCl [pH 7.4], 5 mM MgCl2, 0.5% (v/v) NP-40) containing 1× SIGMAFAST protease inhibitor cocktail (Sigma-Aldrich, S8830). Subsequently, cell debris was pelleted by centrifugation (10 min at 21,910 × g at 4°C) and the supernatant was collected. Protein concentration was determined using Pierce™ 660nm Protein Assay Reagent according to the manufacturer’s instructions (Thermo, 22660). Finally, protein samples were prepared at equal concentrations in 1× Laemmli Loading Buffer (6x: 125 mM Tris-HCl [pH 6.8], 6% (w/v) SDS, 50% (v/v) glycerol, 225 mM dithiotheitol (DTT), 0.1% (w/v) bromophenol blue) and boiled for 5 min. 3D cell cultures were harvested in ice-cold PBS containing 1× SIGMAFAST protease inhibitor cocktail, pelleted by centrifugation at 21,910 × g at 4°C and washed 3 times with PBS containing 1× SIGMAFAST protease inhibitor cocktail to remove the Matrigel. Subsequently, the cell pellets were resuspended in 1× Laemmli Loading Buffer, boiled for 5 min and sonicated.

### Immunoprecipitation

For immunoprecipitations, cells were harvested on ice and lysed by rotation for 1 h at 4°C in JIES buffer (see above) containing 1× SIGMAFAST protease inhibitor cocktail (Sigma-Aldrich, S8830). Subsequently, cell debris was pelleted by centrifugation (10 min at 21,910 × g at 4°C) and the supernatant was incubated with anti-FLAG® M2 Magnetic Beads (Sigma-Aldrich, M8823) or anti-hemagglutinin (HA) agarose beads (Sigma-Aldrich, A2095) overnight at 4°C. The next day, beads were washed 6 times with JIES buffer before the immunoprecipitates were boiled for 5 min in 1× Laemmli Loading Buffer or eluted using 100 μg/mL HA (Thermo Scientific, 26184) or 40 ng/mL FLAG (Generon, A6001) peptide in JIES for 15 min at 1,400 rpm at RT.

### SDS-PAGE and immunoblot analysis

Protein lysates were resolved by sodium dodecyl-sulfate polyacrylamide gel electrophoresis (SDS-PAGE). Protein samples prepared in Laemmli Loading Buffer (see above) were loaded onto either 10%, 12% or 15% (v/v) polyacrylamide gels made in-house, Novex WedgeWell 8-16% gels (Invitrogen, XP08165BOX) or NuPAGE 4-12% Bis-Tris Midi gels (Invitrogen, WG1403A). Electrophoresis was performed in Tris/Glycine/SDS (National diagnostics) or NuPAGE MOPS SDS (Invitrogen, cat. no. NP0001) running buffer. Following SDS-PAGE, the samples were electroblotted onto methanol-activated Amersham Hybond P0.45 polyvinylidene difluoride (PVDF) membranes (GE Healthcare) in Tris/Glycine transfer buffer (National diagnostics). Following transfer, the membrane was blocked with 5% (w/v) milk powder dissolved in PBS/0.1% (v/v) Tween-20 (PBS-T) and incubated with primary antibody for 1 h at RT or overnight at 4°C. Where unconjugated primary antibodies were used, immunoblots were subsequently incubated with horseradish peroxidase (HRP)-conjugated secondary antibodies purchased from Cell Signaling. Signals were visualized using Clarity Western ECL reagent (Bio-Rad) or Femto (Thermo Scientific) using a Vilber Fusion Fx. The following primary antibodies were used: FLAG-HRP (Sigma, A8592, 1:10,000), β-actin-HRP (Abcam, ab49900, 1:25,000), HA-HRP (Roche, 12013819001, 1:1,000-1:5,000), MPP6 (Atlas antibodies, HPA019085, 1:500), RPL27A (Abcam, ab74731, 1:500), LIN7C (Proteintech, 14656-1-AP, 1:1000), MINA (DSHB, 6C6, 1:40), Claudin-1 (Abcam, ab211737, 1:1000).

### Mass spectrometry

MPP6-3xFLAG was anti-FLAG immunopurified from overexpressing HEK293T cells and eluted with 3XFLAG peptide, as outlined above. Samples were extracted, in-solution digested with chymotrypsin, desalted, and analyzed by mass spectrometry as described previously ^29^. Data were searched against the UniProt Reference Homo sapiens database with PEAKS Studio X Software (Bioinformatics Solutions). Cysteine carbamidomethylation was selected as a ‘fixed’ modification, with oxidation (M), deamidation (N, Q) and hydroxylation (P, K, D, N, R, Y, H, W) as variable modifications. The data were then filtered to remove methionine oxidations and those with an A-score of <20 (an A-score of 20 corresponds to a modification correctly localized with 99% certainty).

### *In vitro* hydroxylation assays

For *in vitro* hydroxylation assays, HA-tagged MINA was overexpressed and purified from HEK293T cells, as described above. Following immunoprecipitation, the purified protein was eluted using 2 µg/mL HA peptide (Thermo Scientific, 26184) in 50 mM HEPES, 50 mM NaCl. Purified enzyme was then incubated with RPL27A peptide (GRGNAGGLHHHINFDKYHP) at a final concentration of 100 μM in 50 mM Tris [pH 7.5], 150 mM NaCl, 1 mM DTT for 5 min at 37°C in the presence of 200 μM 2OG, 100 μM Fe(II) and 100 μM ascorbate (final volume 20 μL). For peptide competition assays HA-MINA was incubated with a combination of 50 μM of RPL27A peptide and 50 μM of Rpl8 peptide (NPVEHPFGGGNHQHIGKPSTI) or 50 μM of MPP6 peptide (SQFLEEKRKAFVRRDWDNSGPFCGTISSKKKKK) and incubated for 30 min at 37°C. Thereafter, the reaction was quenched with 10 μL of 1% (v/v) formic acid (HCOOH) in H_2_O. The presence of hydroxylation was subsequently analyzed using a Succinate-Glo JmjC Demethylase/Hydroxylase Assay kit (Promega), as follows. Samples were diluted 1:10 in H_2_O and 5 μL of the diluted reaction was mixed with an equal volume of Succinate-Glo detection reagent 1 and incubated for 1 h at RT in a clear bottom 384-well plate. Each reaction was performed in triplicate. Subsequently, 10 μL of Succinate-Glo detection reagent 2 was added, mixed and incubated for 10 min at RT, before luminescence was analyzed.

### Structural analysis

Molecular modelling and structural comparisons were carried out using the SSM superposition facility ^68^ in Coot ^69^ and visualized in PyMOL (The PyMOL Molecular Graphics System, Version 1.2r3pre, Schrödinger, LLC). Coordinates for human MPP6 full-length were obtained either from the AlphaFold database or generated *ab initio* using the ColabFold server ^70^.

### Statistical analysis

Statistical analyses were performed in R. Statistical significance was determined using unpaired t test or one-way analysis of variance (ANOVA) followed by a Dunnett’s test or pairwise t tests with Benjamini-Hochberg correction for multiple comparisons. Differences were considered significant at p<0.05. Calculated *p* values are shown in each graph. Sample sizes were chosen empirically. Western blot quantifications were performed with Fiji.

## Acknowledgements

We thank Marion Schmidt-Zachmann (DKFZ, Heidelberg) for the MINA antibody and Peter Ratcliffe, Matthew Cockman, and Sally Roberts for critical reading of the manuscript. E.H. was supported by a BBSRC-MIBTP award to E.H. and M.L.C (1790845). E.H, R.A. and U.B. were funded by a Cancer Research UK Programme Foundation Award to M.L.C. (C33483/A2567). S.J.S. is supported by a transitional award from the Francis Crick Institute (CR2019/003DK). The work was also supported by an MRC project grant to M.L.C. (MR/N021053/1).

## Author contributions

E.H. and R.A. designed and performed the cell biology experiments, analyzed the resulting data, and generated the corresponding figures. A.K contributed to generation of data. U.B., J.R.B. and A.Z. contributed to molecular biology. R.H. and R.F. performed mass spectrometry analysis. L.O.-J., R.P., and I.R.P. performed, analyzed, and interpreted multiplex immunofluorescence under the supervision of J.L.Q.. E.H. and M.L.C wrote the manuscript. M.L.C. co-designed the experiments and directed the project. S.J.S. performed structural analyses. All authors approved the final version of the manuscript.

## Declaration of interests

The authors declare no competing interests.

## Notes

### Competing Interest Statement

The authors have declared no competing interest.

### Summary of Updates

Revised Figure 2 to include multiple immunofluorescence of human colon

