## Supplemental Data for "A protein hydroxylase couples epithelial membrane biology to nucleolar ribosome biogenesis"

[1]Institute of Cancer and Genomic Sciences, University of Birmingham, Birmingham, UK
[2]Present address: Department of Biochemistry, Stanford University, Stanford, California, USA
[3]Present address: GSK, Gunnels Wood Rd, Stevenage, UK
[4]Present address: School of Medicine, University of Leeds, Woodhouse, Leeds, UK
[5]CRUK Beatson Institute, Garscube Estate, Glasgow, United Kingdom
[6]Centre for Cellular and Molecular Physiology, University of Oxford, Headington, Oxford, UK
[7]Thermo Fisher Scientific, Cambridge, UK
[8]Target Discovery Institute, Nuffield Department of Medicine, NDM Research Building, University of Oxford, Old Road Campus, Roosevelt Drive, Headington, Oxford, UK
[9]Present address: School of Medicine, UCSF, San Francisco, USA
[10]School of Cancer Sciences, University of Glasgow, Glasgow, United Kingdom.
[11]Department of Histopathology, Queen Elizabeth University Hospital, Glasgow, United Kingdom.
[#]These authors contributed equally

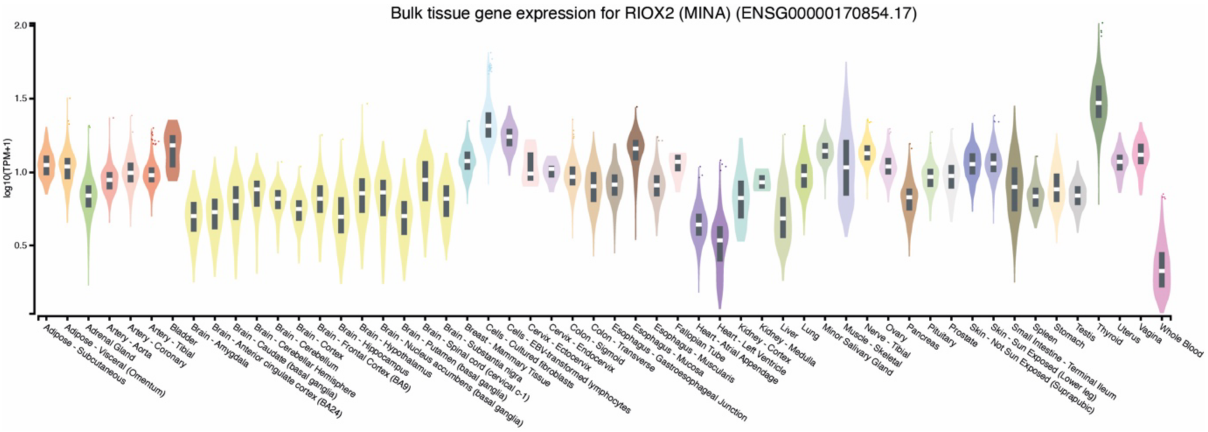


**Figure S1. Ubiquitous expression of MINA in human tissues.** The online database GTEX Portal (version 8) was used to determine the expression of the human RIOX2 (MINA) gene across different tissue types. TPM units are transcripts per million normalized to gene length.

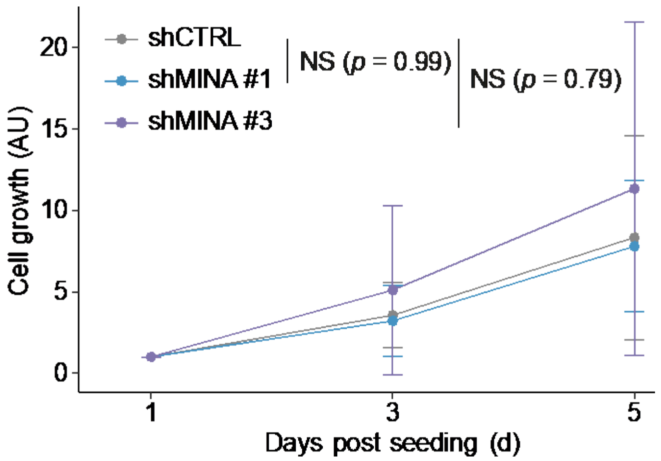


**Figure S2. Loss of MINA does not affect the proliferation of Caco-2 cells.** Cell growth was monitored with resazurin. Data presented are the mean ± SD of $n$ = 4 biological replicates. Four technical replicates of each sample were used in each experiment. No significant differences were observed. AU, arbitrary unit. Statistical significance was determined using a one-way ANOVA and $p$ values were calculated post-hoc using a Dunnett's test.

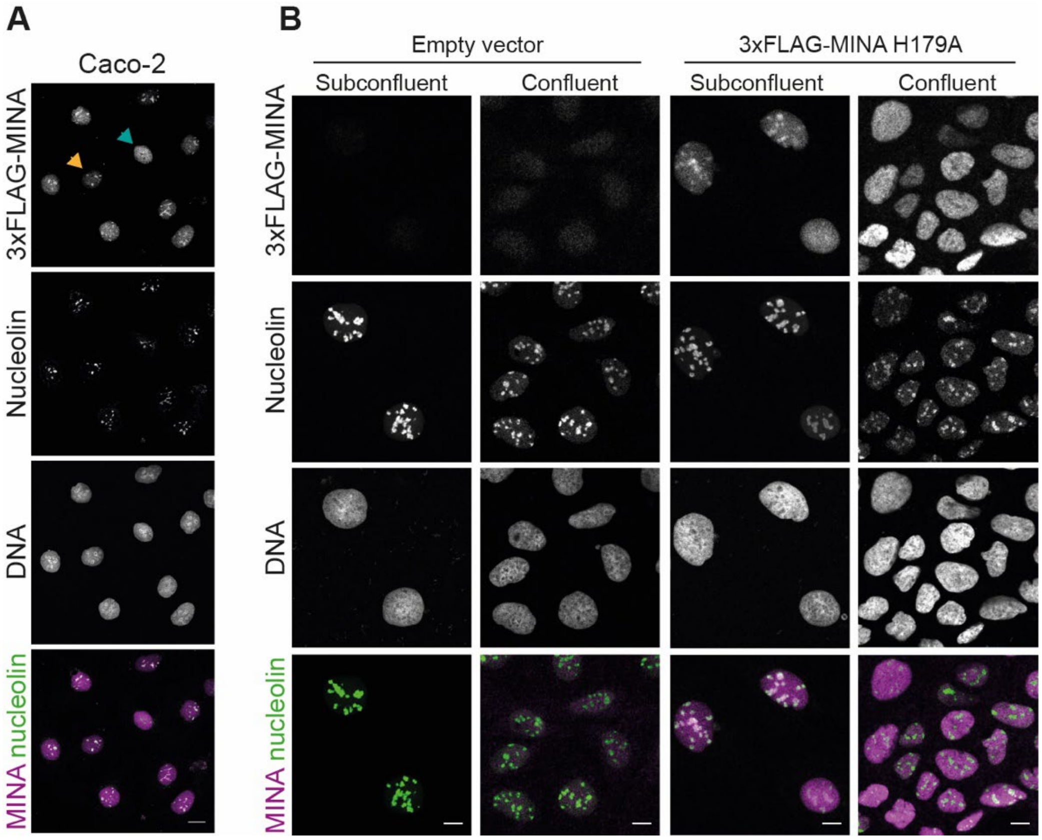


**Figure S3. The nucleolar localization of MINA is regulated by its activity, and by cell growth status.**

**(A)** Differential nucleolar (yellow arrow)/nucleoplasmic (blue arrow) localization of 3xFLAG-MINA in stable doxycycline-inducible Caco-2 cells. Cells were incubated with 0.1 µg/mL doxycycline for 48 h and then fixed with 4% (w/v) PFA and stained for MINA (magenta) and nucleolin (green). Scale bars, 20 µm.

**(B)** Nucleolar enrichment of MINA depends on confluence and catalytic activity**.** Representative immunofluorescence images showing the localization of catalytically inactive (H179A) 3xFLAG-MINA in subconfluent and confluent conditions. Cells were fixed with PFA and stained for MINA (magenta) and the nucleolar marker nucleolin (NCL). Scale bars, 20 µm.

**(A,B)** Nuclei were visualized with DAPI (gray).

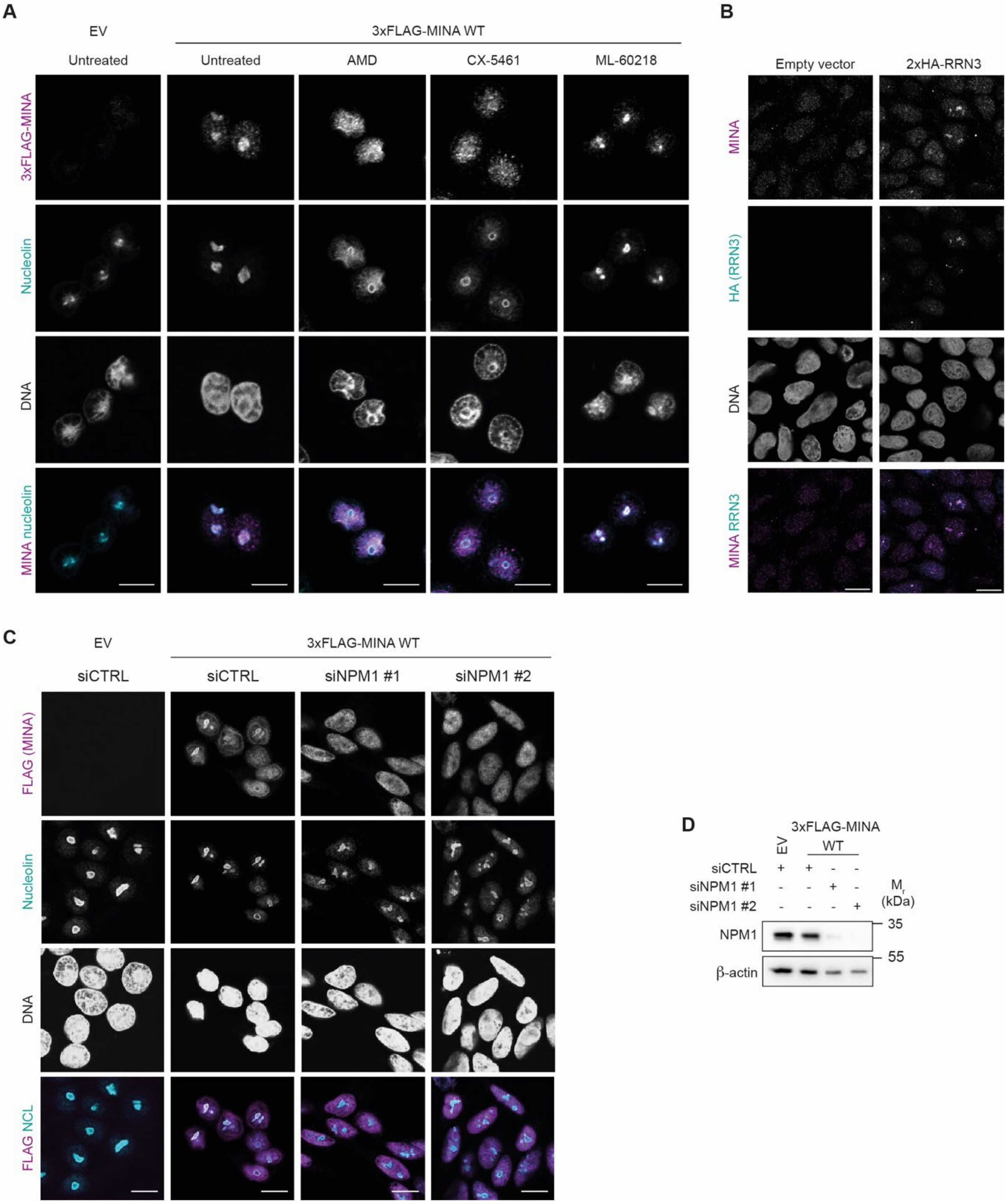


**Figure S4. RNA polymerase I activity and the nucleolar MINA interactome suppress the extra-nucleolar localization of MINA.**

**(A)** Effect of RNA polymerase inhibition on localization of MINA in SW620 cells. Stable SW620 cells expressing an empty vector (EV) or wildtype (WT) 3xFLAG-MINA were treated for 4 h with either DMSO (“Untreated”), RNA polymerase I inhibitors actinomycin D (AMD, 10 ng/mL) or CX-5461 (50 nM), or the Pol III inhibitor ML-60218 (25 μM). Cells were fixed with methanol and stained for MINA (magenta) and nucleolin (NCL, cyan).

**(B)** Nucleolar enrichment of MINA in cells overexpressing the RNA polymerase I transcription factor RRN3. Caco-2 cells stably expressing either an empty vector or doxycycline-inducible 2xHA-RRN3 were grown to a confluent monolayer before expression of 2xHA-RRN3 was induced for 48 h with 1 µg/mL doxycycline. Cells were fixed and stained for MINA (magenta) and HA (cyan). Scale bars, 15 µm

**(C)** Nucleolar enrichment of MINA requires nucleophosmin-1 (NPM1). Stable SW620 cells expressing EV or WT 3xFLAG-MINA were transiently transfected with control (siCTRL) or NPM1-targeting (siNPM1) siRNA. Cells were fixed and stained for FLAG (MINA, magenta) and nucleolin (NCL, cyan).

(**D**) Whole cell extracts from (C) were immunoblotted for NPM1 (33 kDa) and β-actin (42 kDa). Expression of 3xFLAG-MINA was induced with 0.1 µg/mL doxycycline for 48 h.

(**A,B,C**) Nuclei were visualized with DAPI.

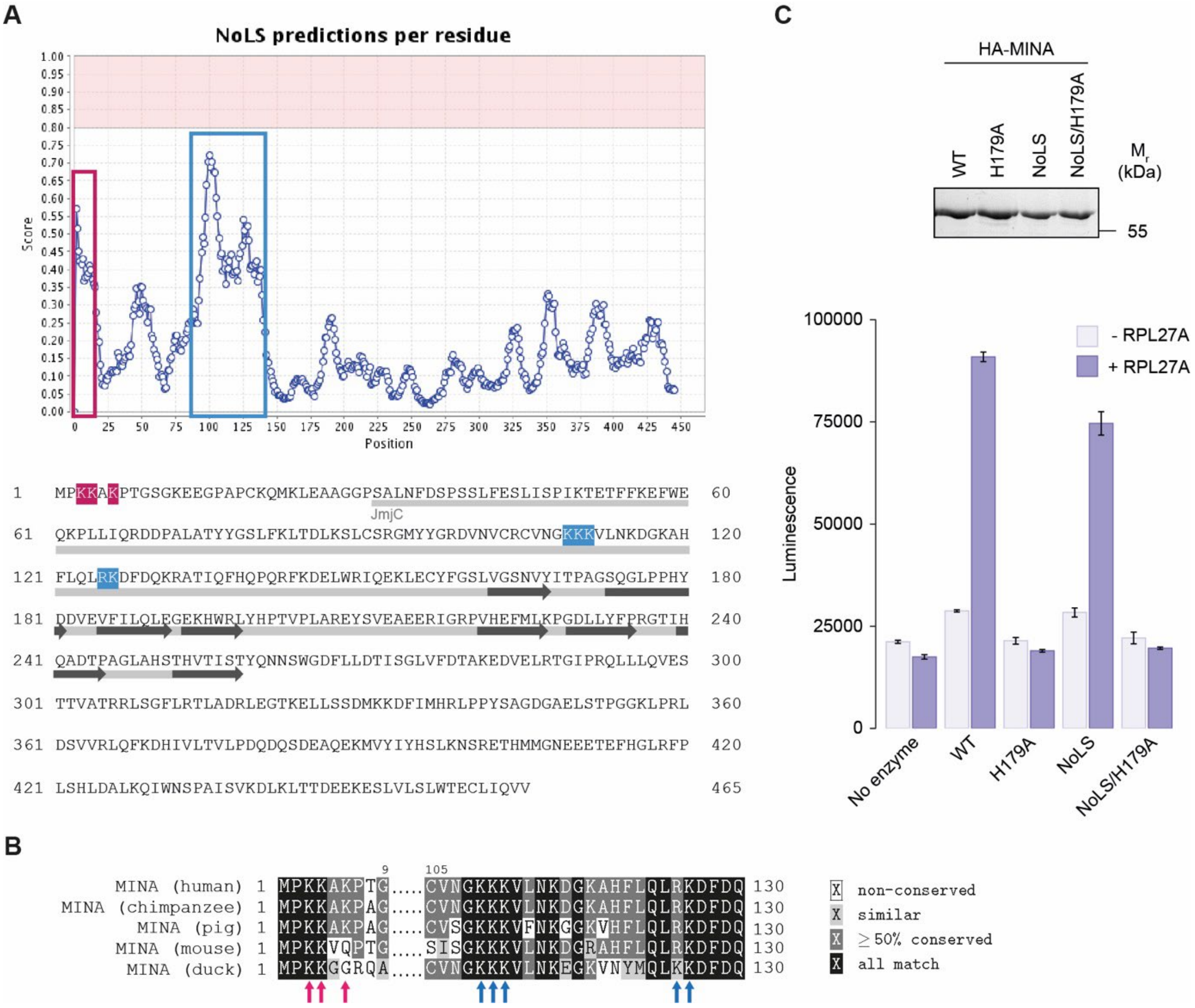


**Figure S5. Identification of primary sequence determinants required for MINA nucleolar localization signal.**

**(A)** Top panel: Nuclear Localization Sequence (NoLS) prediction using online software NoD (http://www.compbio.dundee.ac.uk/www-nod/) identifies two candidate regions in MINA (boxed in pink and blue). Bottom panel: Protein sequence of human MINA. Basic residues contributing to NoD-predicted NoLS are highlighted in pink and blue. The JmjC catalytic domain is underscored in light gray. Dark gray arrows represent the eight β-sheets that make up the catalytic core. UniProt ID = Q8IUF8.

**(B)** Conservation of candidate NoLS residues. Sequence alignment of human (*Homo sapiens*, UniProt ID = Q8IUF8), chimpanzee (*Pan troglodytes*, Uniprot ID = H2QMZ6), pig (*Sus scrofa*, UniProt ID = M3V811), mouse (*Mus musculus*, Q8CD15) and duck (*Anas platyrhynchos platyrhynchos*, Uniprot ID = U3IAG8) MINA protein sequences. Arrows indicate residues of interest within the regions identified as candidate NoLS. Sequence alignment was generated using Clustal Omega.

**(C)** Mutation of R125G/K126G ('NoLS') does not affect the intrinsic enzymatic activity of MINA. *In vitro* hydroxylation assay of RPL27A peptide using wildtype (WT), NoLS or inactive (H179A) HA-MINA immunopurified from transiently transfected HEK293T cells. Top panel: Coomassie blue staining showing the amount of enzyme used in the assay. An uncropped image is presented in source data. Bottom panel: the indicated HA-MINA variants were incubated with 2OG, ascorbate, Fe(II) and with or without RPL27A peptide for 5 min at 37°C before activity was measured indirectly using a Succinate-Glo assay. Result from 1 biological repeat is shown. The experiment was repeated twice with similar results. Values represent the average ± SD of three technical repeats. What appears to be a modest reduction in activity of the NoLS mutant is likely explained by slightly reduced loading in the assay (see coomassie stained gel, above), consistent with reduced expression in HEK293T (e.g. Fig. 2F and Extended Data Fig. 7A). Note that expression of MINA is not negatively impacted by the NoLS mutation in Caco-2 or SW620 cells (e.g. Fig. 3D and Extended Data Fig. 7B, respectively).

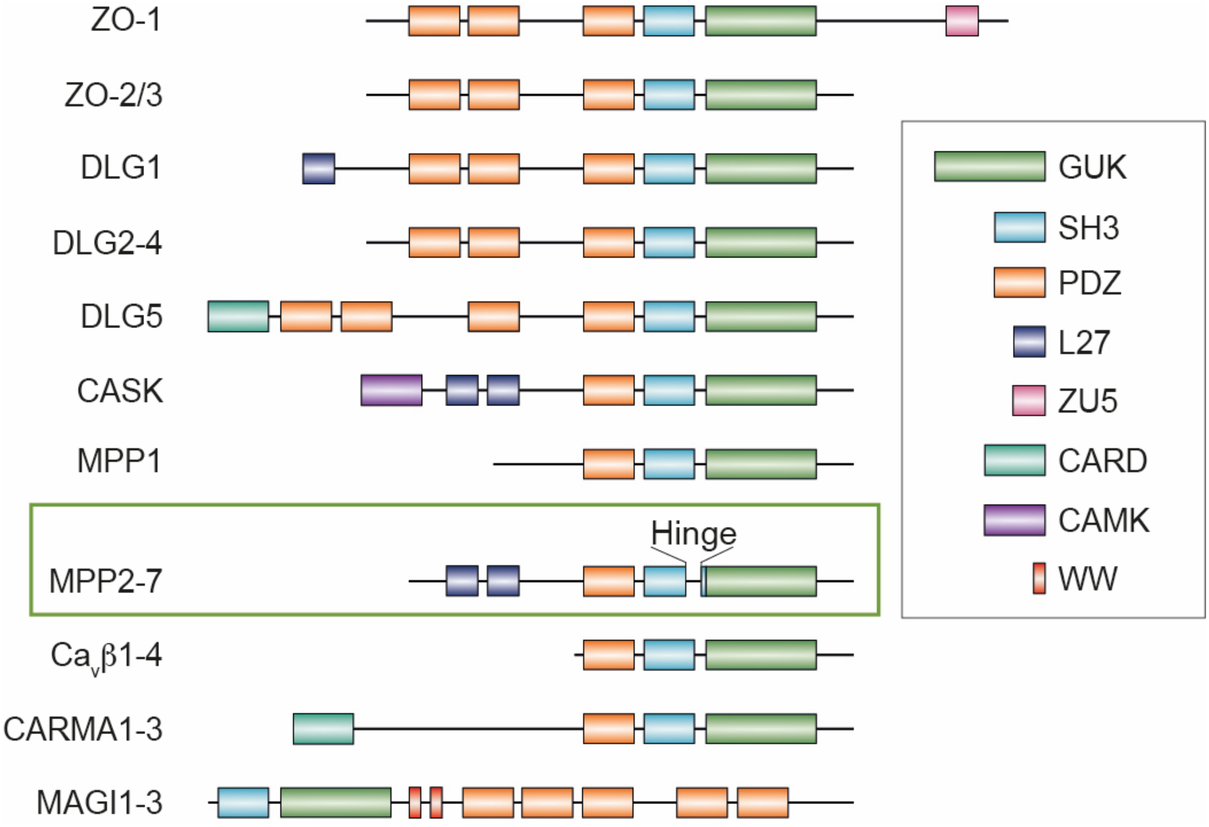


**Figure S6. Domain organization of MAGUK proteins.** With the exception of the Ca$_V$β proteins, all MAGUKs contain (at least one) PSD-95/DLG/ZO-1 (PDZ), src homology 3 (SH3) and guanylate kinase (GUK) domain. The membrane palmitoylated proteins (MPPs), except MPP1, contain two additional L27 domains. The Hinge domain exists as a disordered insert within the SH3 domains, as indicated. We note that the Hinge domain is present in several MAGUKs and is not restricted to MPP2-7. Abbreviations: ZO = zona occludens, DLG = discs large, CASK = calcium/calmodulin-dependent serine protein kinase, Ca$_V$β = voltage-gated calcium channel β subunit, CARMA = caspase recruitment domain-containing MAGUK, MAGI = membrane-associated guanylate kinase inverted.

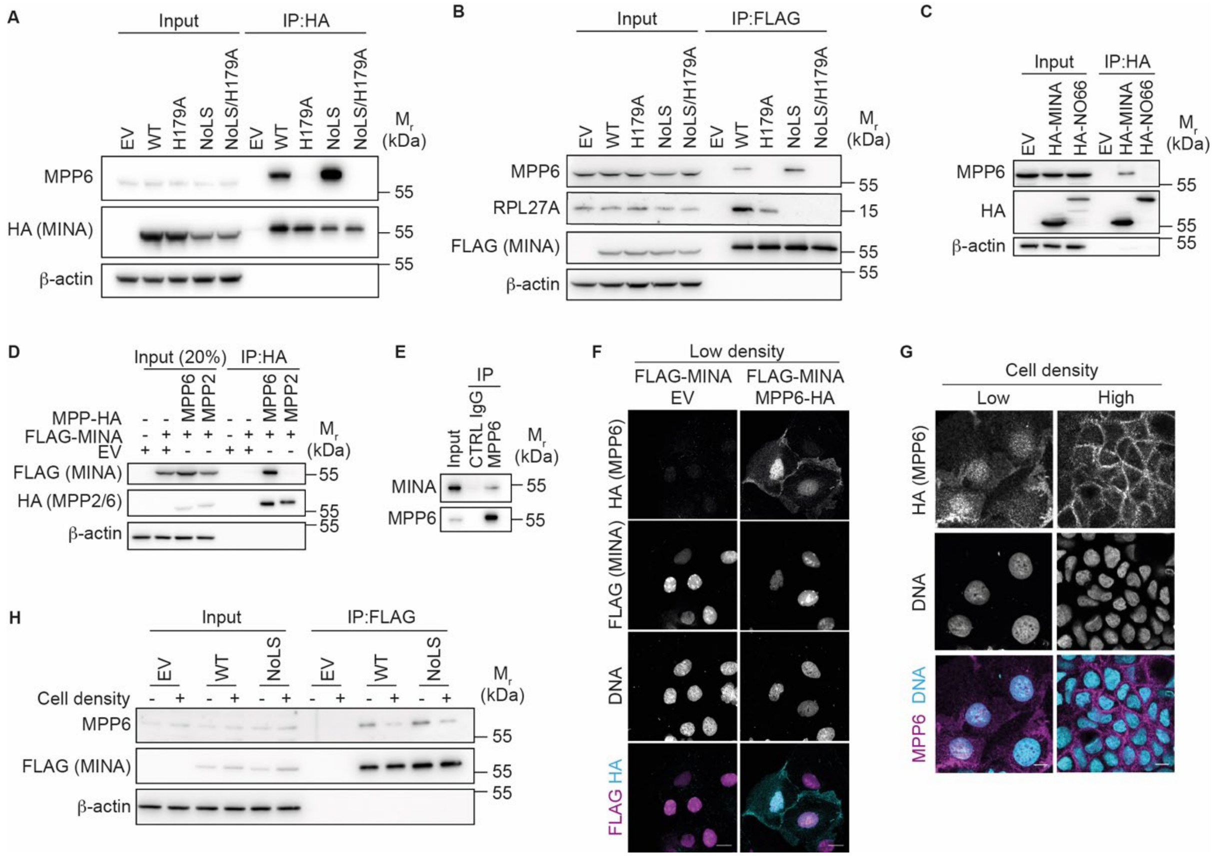


**Figure S7. MINA specifically interacts and colocalizes with MPP6.**

**(A)** Anti-HA immunoprecipitates from extracts of HEK293T cells transiently transfected with an empty vector (EV) or the indicated HA-MINA constructs were immunoblotted for MPP6 (61 kDa) and HA (MINA, 53 kDa). For MPP6 the input is 5% of the IP, for HA and β-actin it is 10%.

**(B)** Anti-FLAG immunoprecipitates from extracts of SW620 cells stably expressing an empty vector (EV) or the indicated doxycycline-inducible 3xFLAG-MINA constructs were immunoblotted for MPP6 (61 kDa), RPL27A (17 kDa) and FLAG (MINA, 53 kDa). Expression of 3xFLAG-MINA was induced using 0.2 µg/mL doxycycline. For MPP6 the input is 2.5% of the IP, for RPL27A it is 0.125%, and for FLAG and β-actin it is 37%.

**(C)** MPP6 binds MINA but not NO66. Anti-HA immunoprecipitates from extracts of HEK293T cells transiently transfected with an EV or vectors encoding HA-MINA or HA-NO66 were immunoblotted for MPP6 (61 kDa) and HA (MINA/NO66). For MPP6 the input is 1.25% of the IP and for HA and β-actin it is 12.5%.

**(D)** MINA binds MPP6 but not MPP2. Anti-HA immunoprecipitates from extracts of HEK293T cells transiently expressing the indicated constructs were immunoblotted for FLAG (MINA, 53 kDa) and HA (MPP2/MPP6).

**(E)** Co-immunoprecipitation of endogenous MINA with MPP6. Endogenous MPP6 was purified from extracts of HEK293T cells and immunoblotted for MINA (53 kDa) and MPP6 (61 kDa).

MINA coprecipitates with an anti-MPP6 antibody but not with the control antibody (CTRL, species-matched control IgG) IP. For MPP6 the input is 11% of the IP and for MINA it is 0.14%.

**(F)** Co-localization of 3xFLAG-MINA and MPP6-HA in the nuclei of Caco-2 cells under low density culture conditions. Caco-2 cells stably expressing an EV or doxycycline-inducible 3xFLAG-MINA were transiently transfected with MPP6-HA, fixed with PFA and stained for HA (MPP6, cyan) and FLAG (MINA, magenta). Scale bars, 20 μm. Expression of 3xFLAG-MINA was induced using 0.1 μg/mL doxycycline for 48 h.

**(G)** MPP6-HA localizes to the nucleus in low density Caco-2 cells. Stable Caco-2 cells constitutively expressing MPP6-HA were seeded at low or high density, fixed with PFA and stained for HA (magenta). Scale bars, 10 μm.

**(H)** MINA and MPP6 preferentially interact in subconfluent cells. Anti-FLAG immunoprecipitates from extracts of low (-) or high (+) density stable Caco-2 cells expressing the indicated constructs were immunoblotted for MPP6 (61 kDa) and FLAG (MINA, 53kDa). Expression of 3xFLAG-MINA was induced for 48 h using 1 μg/mL doxycycline. For MPP6 the input is 0.25% of the IP and for FLAG and β-actin it is 0.8%.

(F) and (G) Nuclei were visualized with DAPI (cyan). (A) and (D) to (H); β-actin (42 kDa) was used as loading control. Uncropped western blot images are presented in source data.

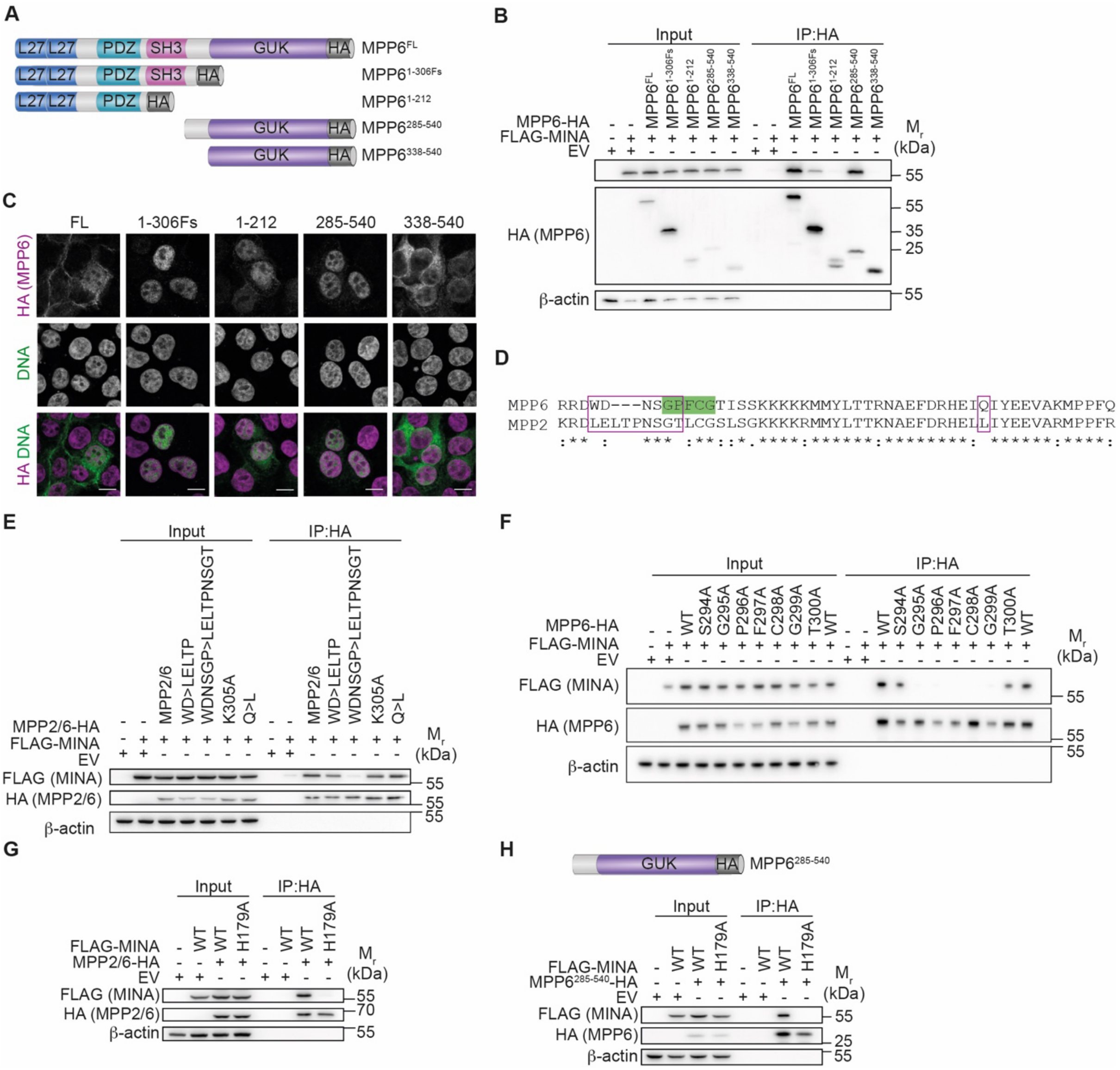


**Figure S8. The interaction with MINA is mediated by specific residues within the Hinge domain of MPP6.**

**(A)** Domain structures of MPP6 mutants used in (B,C). FL = full-length, 306Fs = a variant with a frameshift mutation at amino acid 306.

**(B)** MPP6 amino acids 285-338 are required for coprecipitation of FLAG-MINA**.** Anti-HA immunoprecipitates from extracts of HEK293T cells transiently expressing an empty vector (EV) or the indicated FLAG-MINA and MPP6-HA constructs were immunoblotted for FLAG (MINA, 53 kDa) and HA (MPP6, 61 kDa). For HA the input is 20% of the IP and for FLAG and β-actin it is 10%.

**(C)** Localization of MPP6 truncation mutants, demonstrating their availability for MINA binding in the nucleus. HEK293T cells transiently expressing the indicated MPP6-HA constructs were fixed with PFA and stained for HA (MPP6, green). Nuclei were visualized with DAPI (magenta). Scale bars, 10 $\mu$m.

**(D)** Alignment of human MPP6 (Uniprot ID = Q9NZW5) amino acids 285-338 with MPP2 (Uniprot ID = Q14168). Sequence alignment was generated using Clustal Omega. Note the lack of conservation in the boxed areas, which were the focus of the interaction assays that follow.

**(E)** The LELTPNSGT sequence prevents MPP2 from binding MINA. The indicated changes were made in the context of the MPP2/6 chimera (Fig. 4F). Anti-HA immunoprecipitates from extracts of HEK293T cells transiently expressing the indicated constructs were immunoblotted for FLAG (MINA, 53 kDa) and HA (MPP2/6). Input = 50% (FLAG, $\beta$-actin), 40% (HA).

**(F)** AAs 295-299 are required for MINA binding. Substitution of G295, P296, F298, C298 or G299 to alanine interrupts MINA binding. Anti-HA immunoprecipitates from extracts of HEK293T cells transiently expressing the indicated constructs were immunoblotted for FLAG (MINA, 53 kDa) and HA (MPP6, 61 kDa). Input = 20% (HA), 10% (FLAG).

**(G)** MPP2/6 interacts with MINA in an activity-dependent manner. Anti-HA immunoprecipitates from extracts of HEK293T cells transiently expressing the indicated constructs were immunoblotted for FLAG (MINA, 53 kDa) and HA. Input = 10% (FLAG, $\beta$-actin), 100% (HA).

**(H)** $MPP6^{285-540}$ interacts with MINA in an activity-dependent manner. Anti-HA immunoprecipitates from extracts of HEK293T cells transiently expressing the indicated constructs were immunoblotted for FLAG and HA. Input = 10% (FLAG, $\beta$-actin), 20% (HA). (B), (E), and (H); $\beta$-actin (42 kDa) was used as loading control. EV = empty vector. Uncropped western blot images are presented in source data.

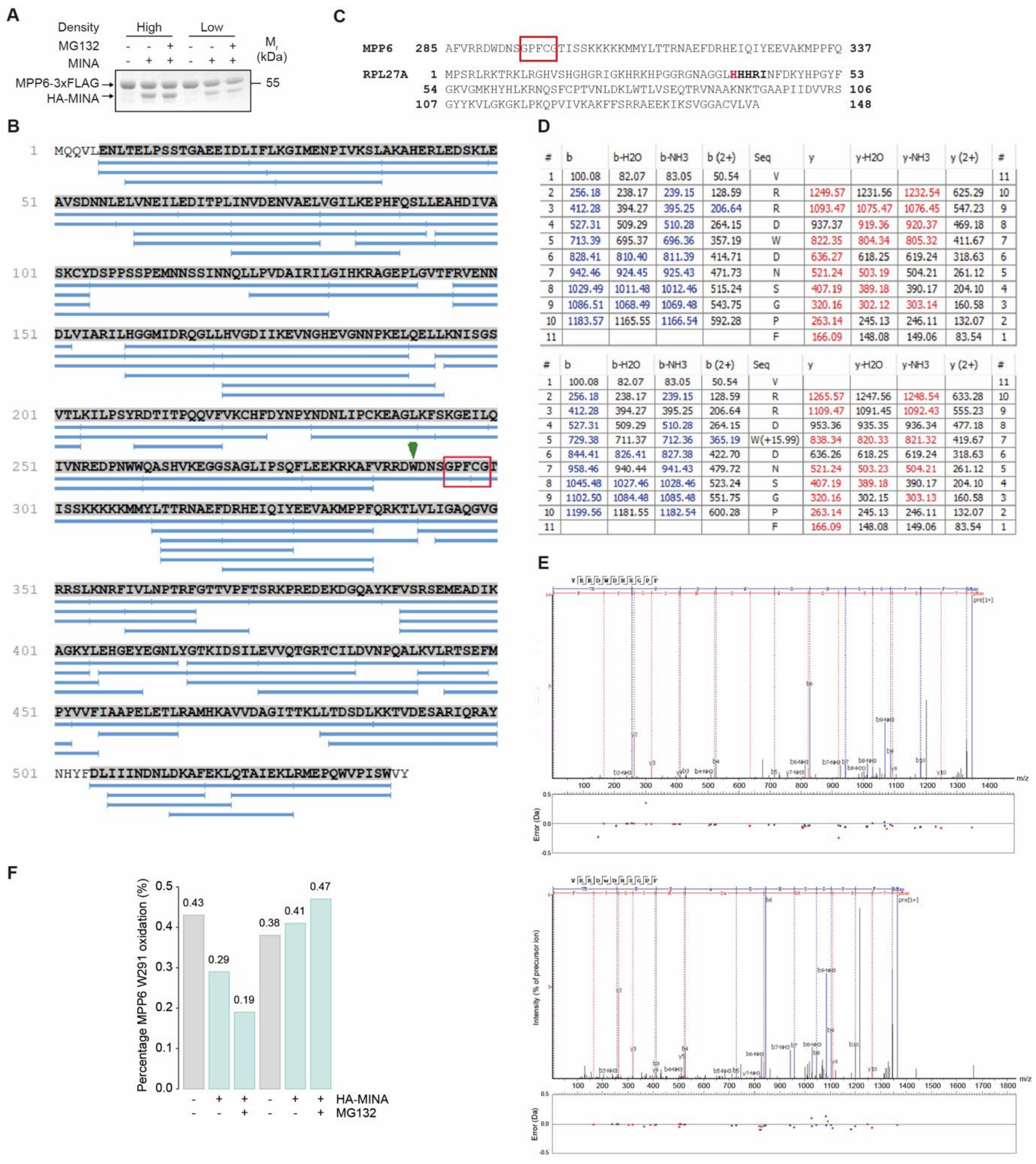


**Figure S9. MPP6 W291 oxidation is low stoichiometry and does not increase in response to MINA overexpression, proteasome inhibition or confluence.**

**(A)** Coomassie Brilliant Blue staining of samples submitted for LC-MS/MS analysis shows efficient purification of MPP6 (upper band) and co-precipitation of MINA (lower band). An uncropped image is presented in source data.

**(B)** Chymotryptic digest coverage (98%) of human MPP6. Blue bars represent identified peptides at a False Discovery Rate of 2%. The minimal MINA binding domain is indicated by a red box. We detected a high scoring (A-score 1000) oxidized tryptophan (W291) (green arrow) in close proximity to the MINA binding domain. An A-score of 20 corresponds to a modification correctly localized with 99% certainty.

**(C)** Comparison of the HINGE/Hook domain of MPP6 containing the minimal MINA binding domain (red box) to the MINA substrate RPL27A (target site bolded, H39 hydroxylation site in red).

**(D)** Tables listing MS/MS fragments and ions identified from the MPP6 chymotryptic non-oxidized (upper) and typtophan-oxidized (lower) VRRD**W**DNSGPF peptides showing theoretical (black) and identified (blue/red) ion species.

**(E)** MS/MS spectra of non-oxidized (upper) and typtophan-oxidized (lower) VRRD**W**DNSGPF peptides.

**(F)** Bars indicate the percentage of MPP6 oxidation at W291 in the various samples indicated, as determined by LC-MS quantification. Note the very low stoichiometry (<0.5%) and the lack of positive response to the indicated treatments.

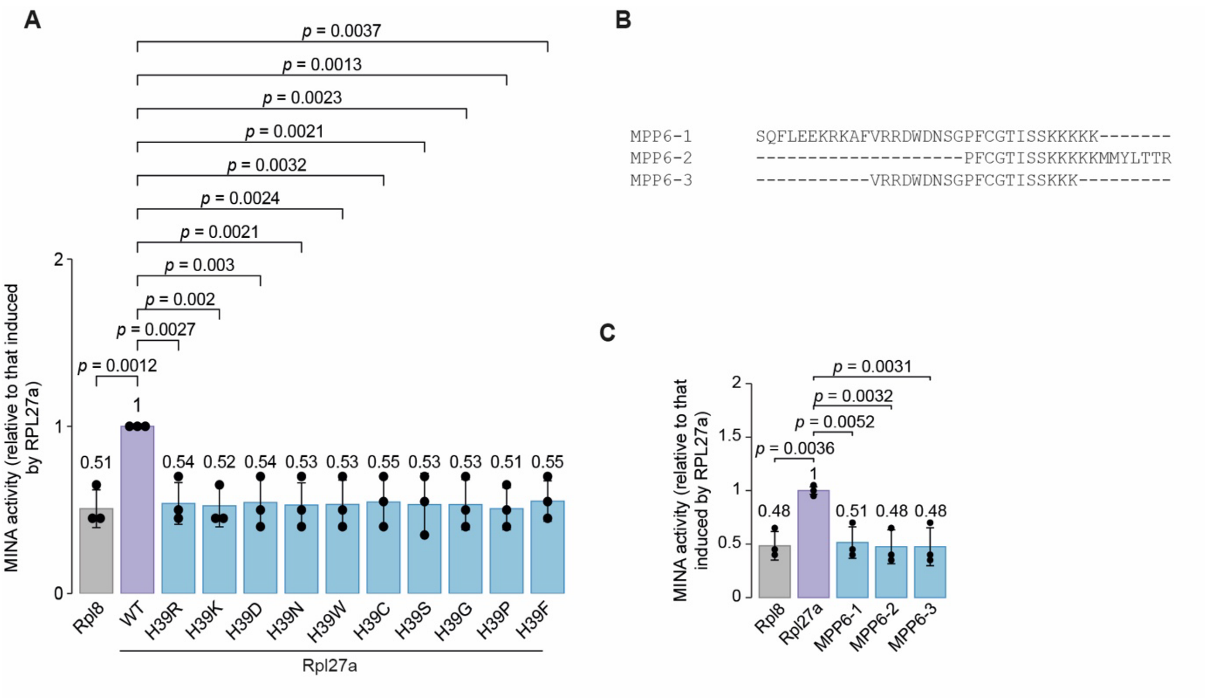


**Figure S10. In vitro enzymatic assays do not support hydroxylation activity of MINA towards MPP6.**

**(A)** Substitution of His-39 to Arg (H39R), Lys (H39K), Asp (H39D), Asn (H39N), Trp (H39W), Cys (H39C), Ser (H39S), Gly (H39G), Pro (H39P) or Phe (H39F) does not support hydroxylation of RPL27A peptide catalyzed by MINA, unlike the wildtype (WT) peptide.

**(B)** Amino acid sequence of the MPP6 peptides used in (B)**.**

**(C)** MINA does not support hydroxylation of MPP6 peptides in vitro.

(A) and (C) Bars indicate the average of n = 3 biological repeats. Averages are presented above each respective bar. Error bars, SD. Statistical significance was determined using a one-way ANOVA and *p* values were calculated post-hoc using Dunnett's multiple comparison test. NO66 substrate Rpl8 was used as negative control. Statistical source data are provided.

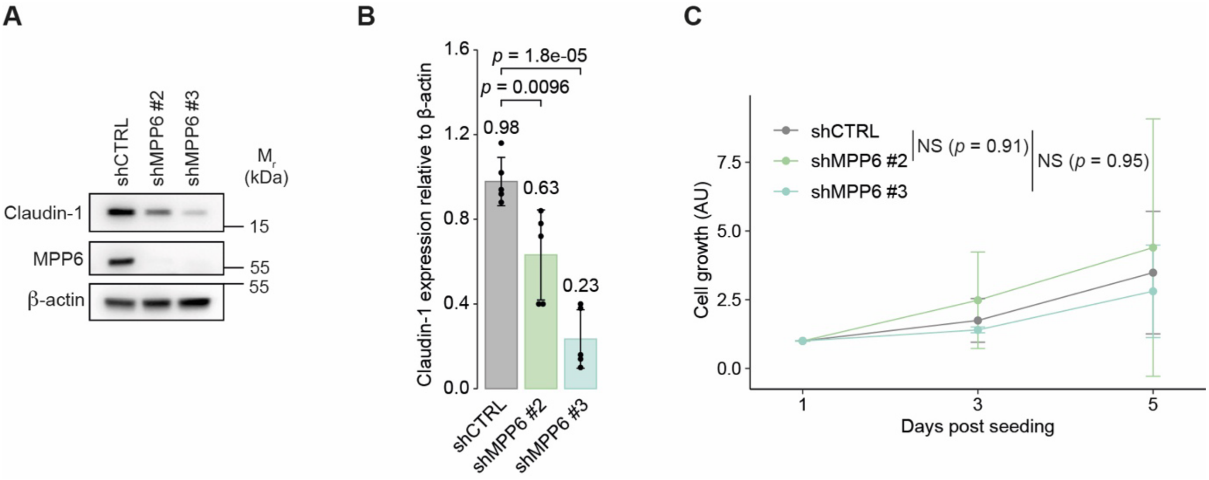


**Figure S11. MPP6 promotes epithelial barrier function without affecting proliferation.**

**(A)** Reduced Claudin-1 expression in response to MPP6 depletion. Extracts from Caco-2 cells expressing the indicated shRNA sequences were grown to a monolayer for 5 days and immunoblotted for Claudin-1 (23 kDa) and MPP6 (61 kDa). β-actin was used as loading control. Uncropped images are shown in source data.

**(B)** Quantification of (A). Bar graph shows Claudin-1 expression relative to β-actin. Bars represent the average of $n$ = 5 biological repeats.

(**C**) MPP6 depletion does not affect cell proliferation. Cell growth was measured in the indicated days with resazurin. Lines represent the average of n = 3 independent biological repeats.

(A and C) Statistical significance was determined using a one-way ANOVA and $p$ values were calculated post-hoc using a Dunnett's test. Error bars, SD.

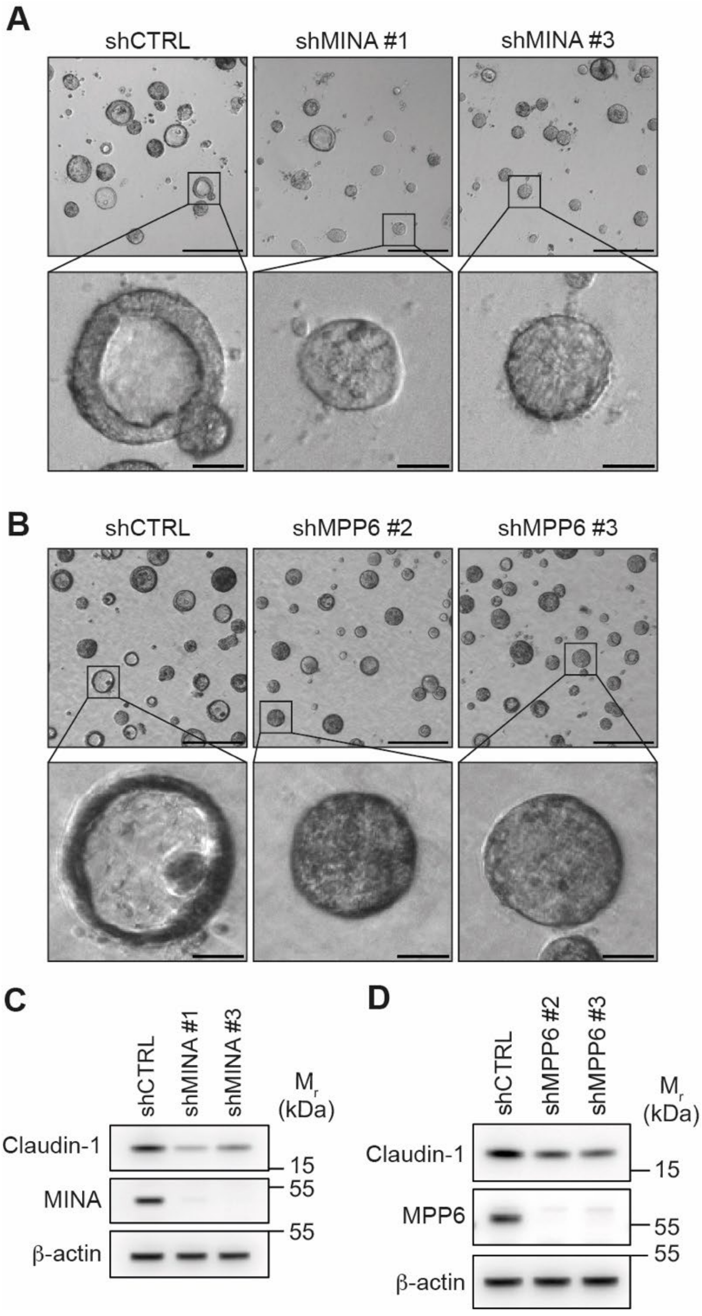


**Figure S12. The MINA-MPP6 pathway is required for lumenogenesis, growth and polarity.**

**(A and B)** Phase-contrast images of control, MINA or MPP6 shRNA-expressing spheroids, grown for 7 days in matrigel. Scale bars, 250 or 30 (enlargements) μm.

**(C and D)** Reduced Claudin-1 expression in MINA (C) or MPP6 (D) shRNA-expressing spheroids. Whole cell extracts of the indicated spheroids were immunoblotted for Claudin-1 (23 kDa) and MINA (C; 53 kDa) or MPP6 (D; 61 kDa). β-actin (42 kDa) was used as loading control.

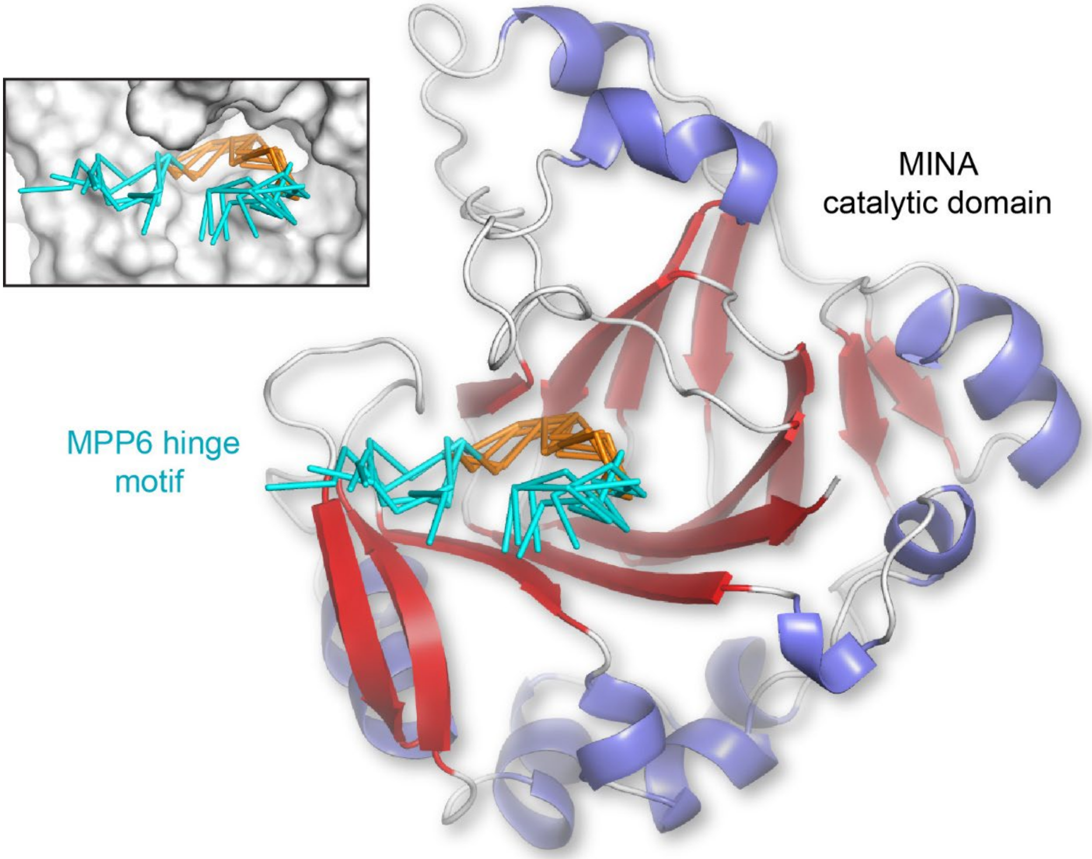


**Figure S13. AlphaFold modelling of the interaction of the MPP6 Hinge motif with the catalytic domain of MINA.** The catalytic domain of MINA is shown in a cartoon representation ($\alpha$-helices in blue, $\beta$-sheets in red) along with five predicted conformations of the MPP6 hinge segment shown as a C$\alpha$ trace with the GPFCG motif coloured orange. Potential interaction with the active site cleft is highlighted by the surface representation shown in the inset.
